# A Stable Nano-Vaccine for the Targeted Delivery of Tumor-Associated Glycopeptide Antigens

**DOI:** 10.1101/2021.04.27.438445

**Authors:** Kevin R. Trabbic, Kristopher A. Kleski, Joseph J Barchi

**Author notes:** Address Correspondence to: Joseph J Barchi Jr, Head, Glycoconjugate and NMR Section, Chemical Biology Laboratory, Center for Cancer Research, 376 Boyles Street, Bldg 376, Rm 209, NCI Frederick, Frederick, MD 21702, (Voice).

## Abstract

We have developed a novel antigen delivery system based on polysaccharide-coated gold nanoparticles (AuNPs) targeted to antigen presenting cells (APCs) expressing Dectin-1. AuNPs were synthesized de-novo using yeast-derived β-1,3-glucans (B13Gs) as the reductant and passivating agent in a microwave-catalyzed procedure yielding highly uniform and serum-stable particles. These were further functionalized with both peptides and glycopeptides from the tandem repeat sequence of mucin 4 (MUC4), a glycoprotein overexpressed in pancreatic tumors. The glycosylated sequence contained the Thomsen-Friedenreich disaccharide, a pan-carcinoma, Tumor-Associated Carbohydrate Antigen (TACA), which has been a traditional target for antitumor vaccine design. These motifs were prepared with a cathepsin B protease cleavage site (Gly-Phe-Leu-Gly), loaded on the B13Gs-coated particles and these constructs were examined for Dectin-1 binding, APC processing and presentation in a model in vitro system and for immune responses in mice. We showed that these particles elicit strong in vivo immune responses through the production of both high-titer antibodies and priming of antigen-recognizing T-cells. Further examination showed that a favorable antitumor balance of expressed cytokines was generated, with limited expression of immunosuppressive Il-10. This system is modular in that any range of antigens can be conjugated to our particles and efficiently delivered to APCs expressing Dectin-1.

## INTRODUCTION

Tumor-associated carbohydrate antigens (TACAs) are glycan structures covalently attached to proteins in various forms on the surface of tumor cells.^1–3^ They differ from the normal cell glycan repertoire insofar as the tumor biosynthetic machinery is modified via a disparate regulation of glycosyltransferases and hydrolases. This produces aberrant and distinct cellsurface glycan structures that are unique to tumors, and these structures impart modified biophysical and protein binding characteristics to individual tumor types. In addition, some of these tumor-associated glycans can be recognized as “foreign” by the immune system (hence the moniker, “antigen”) eliciting both humoral and (sometimes) cell-mediated responses. As a result, there have been myriad attempts to prepare vaccine constructs to raise effective and durable immune responses to TACAs.^4–13^

TACAs are composed of self-molecules, and hence are innately weak immunogens; this makes them “tolerant” to robust immune responses. Carbohydrates are also T-cell independent epitopes and mobilization of that arm of the immune response is essential to eradicate any established tumor.^10^ It is thus not surprising that vaccine development against these antigens has been problematic; however, many strategies have met with success. The use of various delivery platforms^4, 14–17^, conjugations to immunogenic proteins (KLH, Tetanus toxin)^18^ or peptide epitopes to generate T-cell help^19–22^, covalent conjugation to Toll-like receptor agonists,^6, 18, 20, 23^ the use of various adjuvants^24^ and attachment to zwitterionic polysaccharides^25, 26^ have led to robust immune responses to carbohydrate structures and some have moved into clinical trials. However, none of these efforts have led to a truly effective, FDA approved therapy against any type of cancer.

Research into the use of different nanomaterials for a host of medical applications has exploded in recent years. Not surprisingly, many of these applications are for some form of targeted drug delivery to treat various ailments in animal models. It follows that some of these same materials have been purposed as vaccine platforms to deliver antigens, adjuvant and T-cell epitopes, either alone or in some combinations to generate functional immune responses against disease.^27^ Along with applications to infectious diseases, the search for anticancer vaccines has spawned much work in this area. The list of nanomaterials employed is now quite extensive, and those include metallic nanoparticles, glycodendrimers, liposomes and natural materials.^14, 28^ Some examples of TACA/nanoparticle-based platforms for cancer immunotherapy include, 1) liposome and lipid-based particles,^29^ 2) Metallic, and/or ferromagnetic particles^30–32^, 3) synthetic biodegradable polymers or hydrogels^33^ and 4) Virus-Like Particles (VLPs).^17, 34, 35^ Of the metallic particles, gold nanoparticles (AuNPs) have emerged as the most versatile and hence most utilized for various therapeutic applications. They can be easily synthesized in a size selective manner and be coated with most any appropriately functionalized molecular family (proteins, small molecules, carbohydrates, lipids). These features led us to develop various AuNPs as either antitumor therapies or vaccines.^36–41^ Our original vaccine platform utilized a complex mixture of a glycopeptide antigen and molecular adjuvant all coated on a gold nanoparticle where we added a spacer to reduce antigen density.^37^ The glycopeptide was derived from the tandem repeat unit of a mucin protein (MUC4) that is over/aberrantly expressed on pancreatic ductal adenocarcinomas,^42^ and contained glycosylated serine or threonine residues containing the Thomsen Friedenreich (TF) antigen, a well-known TACA presented almost exclusively on tumor tissue.^43, 44^ This first generation produced a defined but weak antibody response with little cell-mediated response. We set out to redefine the design with a newly constructed particle that could be targeted to Antigen Presenting Cells (APC’s), such as macrophages and dendritic cells, but be versatile enough to allow further modifications with appropriate components to facilitate robust immune responses.

APC’s present a wide variety of receptors on their surface that facilitate binding and uptake of foreign antigens displayed on bacteria, viruses, fungi and tumors. These are sometime referred to as pattern recognition receptors (PRRs) as they recognize specific foreign antigens on microbes (so called Pathogen-Associated Molecular Patterns, PAMPs) as part of the innate immune response.^45^ Two categories of PRRs are the Toll-like receptors (TLRs)^46, 47^ and C-type lectins (CTLs), or calcium-dependent carbohydrate binding proteins (CBPs).^48^ These specialized proteins all recognize distinct molecular patterns, each driving various cellular signaling events that lead to cytokine production, immune cell activation and mobilization. Both sets of receptors have been targeted to stimulate responses in antimicrobial or antitumor immunotherapy. Many CTLs have been targeted through conjugation of their cognate carbohydrate ligands onto various platforms as therapeutic strategies against infectious diseases and cancer. Some CTLs that have been targeted include Dendritic Cell-Specific Intercellular adhesion molecule-3-Grabbing Non-integrin (DC-SIGN)^15, 49^, the Mannose Receptor (MR)^15^, Macrophage-Galactose type C-type Lectin (MGL)^50–53^ and the Dectins (1, 2 and 3).^54–56^ Dectin-1 is a CTL that binds β-1,3-glucans (B13Gs), the most abundant polysaccharide in many fungal species; this engagement initiates signaling which is mediated through an intracellular immune-receptor tyrosine-based activation motif (ITAM).^48^ Tyrosine phosphorylation by Src-family kinases initiates a signaling cascade leading to NF-kB activation and the production of various inflammatory cytokines.^48^ In addition, targeting dectin-1 with B13Gs has been a strategy to deliver antigens to APCs, as engagement with B13Gs initiates endocytosis leading to antigen presentation via MHC-II molecules.^57^ B13Gs have been widely used as immune stimulants for their ability to kick-start the production of reactive oxygen species (ROS,) inflammatory cytokines and microbial killing. The dectin-1/B13Gs signaling system has been referred to as a bridge between innate and adaptive immunity.^58^

In this work we combined the strategy of using AuNPs as a delivery platform, but designed in such a way to target Dectin-1 on APCs via B13Gs; this would deliver glycopeptide antigens derived from tumor-associated mucins (*vide supra*) in an effort to generate a glycopeptide-specific immune response. The B13G polysaccharide was used as both the reductant and the stabilizing agent to create a platform that could further be coated with antigenic sequences. There have been several studies that have prepared AuNPs in this way using various naturally-derived gums and other saccharide components^59–62^, as well as B13Gs.^63, 64^ However, none of these studies ustilized the polysaccharide for as a targeting agent. Here we report a simple synthetic strategy toward these multifunctional AuNPs to deliver glycopeptide antigens and raise potent immune response to these antigens as measured by antibody and cytokine production.

## RESULTS AND DISCUSSION

We were encouraged from our previous study^37^ to pursue a refined AuNP platform for a vaccine that can deliver glycopeptide antigens derived from proteins that are overexpressed in tumors that display various covalently linked TACAs. There are a variety of carriers that have been studied, many based on inorganic nanoparticles or some type of “organic” nano-construction, e.g. from modified viruses (modified adenoviruses, bacteriophage and virus capsid proteins). However, due to the simplicity of AuNPs synthesis and manipulation it was decided to develop a modular AuNP system that could be delivered to APC’s easily and efficiently. We chose the Dectin-1/β-1,3-glucan system based on the following: 1) Dectin-1 is an atypical CLR in that is does not have a requirement for calcium^65^, and so activity would not be as sensitive to divalent metals; 2) Signaling through Dectin-1/β-1,3-glucan elicits the production of inflammatory cytokines that may skew the T-cell repertoire toward and antitumor response^66^ and 3) Many polysaccharides and gums have been used to prepare AuNPs^61, 67–69^, as well as B13Gs, but they have not been utilized for TACA-containing glycopeptide antigen delivery. A schematic representation of our overall plan is shown in Figure 1.

**Figure 1.**
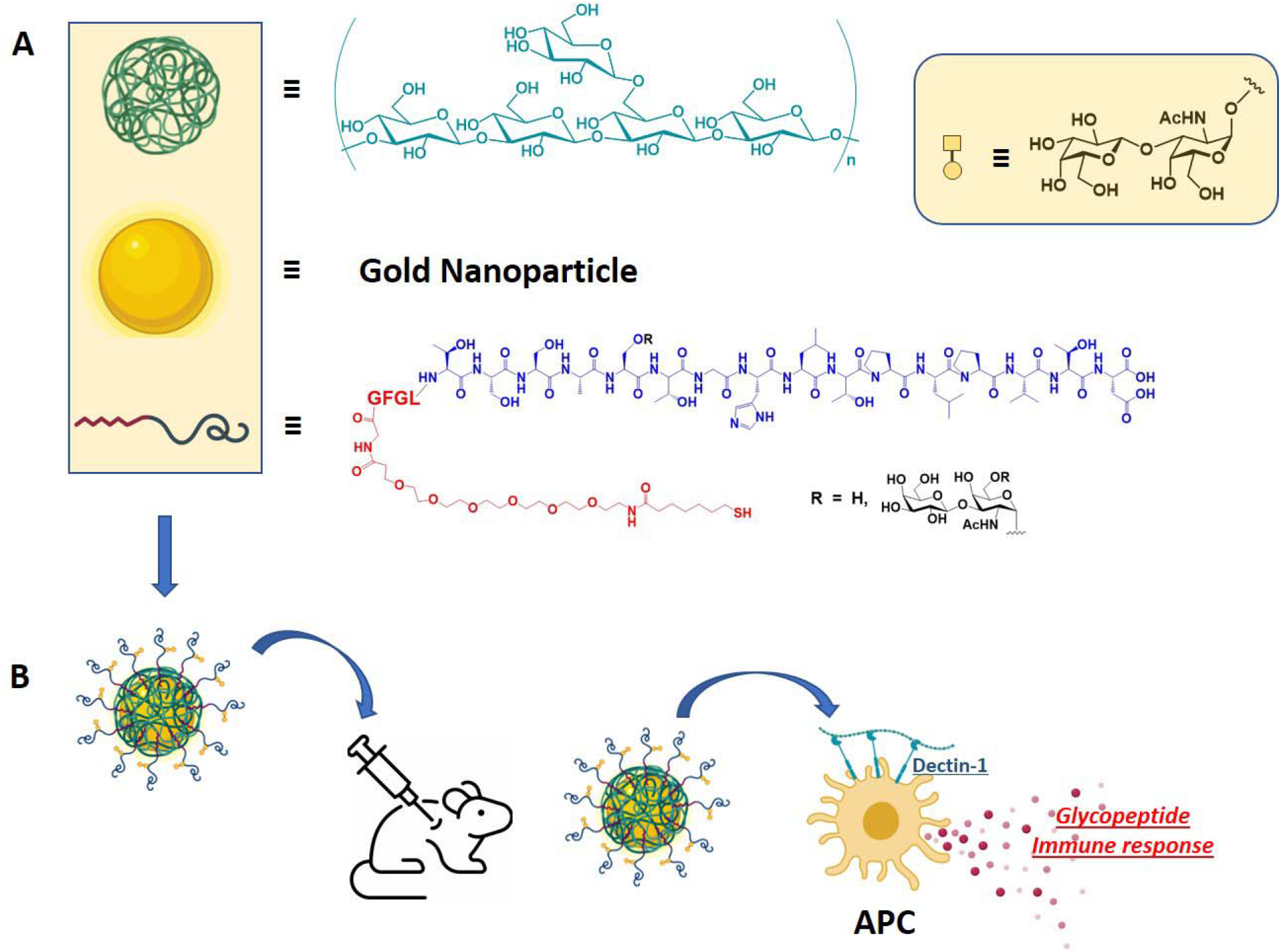
Schematic representation of particle components (A); β-1,3-glucan (green), gold nanoparticle (gold) and MUC4 glycopeptide antigen (red/dark blue curved line; red = linker, blue = glycopeptide); tetrapeptide -GFGL- was inserted as a cathepsin B cleavage site. (B) Combined nanoparticle used for in vivo production of specific immune responses in mice.

### Peptide/Glycopeptide/Linker Synthesis

The synthesis of the various peptides and glycopeptides followed from our previous studies with slight modifications.^37, 41^ The linker was the same we used in our vaccine work since we had shown then and more recently^70^ that this was a robust and non-immunogenic motif to connect our antigens to nanoparticles. For this work, we used a CEM^®^ PRIME Microwave peptide synthesizer to prepare all peptides. This instrument uses Oxyma/Diisopropylcarbodiimide activation with 2 min peptide coupling cycles at 90°C. Most glycoamino acid couplings were performed manually using conditions developed previously to minimize α-carbon amino acid epimerization.^71^ Those performed on the PRIME instrument included an equivalent of Hunig’s base (diisopropylethyl amine, DIEA) to offset the slightly acidic conditions afforded in Oxyma-activated amino acid couplings. Deprotection of the carbohydrate acetate groups was affected by treatment with 0.5 M sodium methoxide in methanol solution. All peptides were purified by reverse phase HPLC and characterized by ESI and MALDI mass spectrometry, as well as NMR spectroscopy (Supporting Information)

### Synthesis and Characterization of B13Gs-Coated AuNPs

There is precedent for the de-novo preparation of AuNPs employing the reducing end of a polysaccharide to both reduce Au^+3^ to Au^0^ with simultaneous passivation of the resulting particles.^61^ As for other AuNP syntheses, relative concentration, temperature and reaction conditions will dictate the size and quality of the particles. The physical characteristics of the polysaccharide offer challenges, such as solubility and issues with interconverting tertiary structures and conformations. B13Gs are subject to these challenges; they assume triple helix structures in solution and often need high pH to denature and hence solubilize the polymers.^72–77^ Our synthesis started with the dissolution of the B13Gs in 1M NaOH solution with heating under microwave irradiation. We found that use of microwaves to be essential to the efficient and high-quality synthesis of the nanoparticles. After dissolution in base, the AuNPs form smoothly in about 90 min under microwave irradiation. The synthesis and select characterization data for B13G AuNPs and those coated with OVA peptides (as part of our in vitro model study, *vide infra*) are shown in Figure 2. The particles are very uniform by Transmission Electron Microscopy (TEM, Figure 2A,B) and Dynamic Light Scattering (DLS, Figure 2C,D), with average core diameters of 15-17 nm and hydrodynamic diameters much larger at ~40 nm, indicative of a large, and possibly highly hydrated polymer coated on the particle surface.^78^ We found the procedure to be very reproducible and the batch-to-batch size measurements to be highly comparable. (see Supporting Information Figures S1-S4)

**Figure 2.**
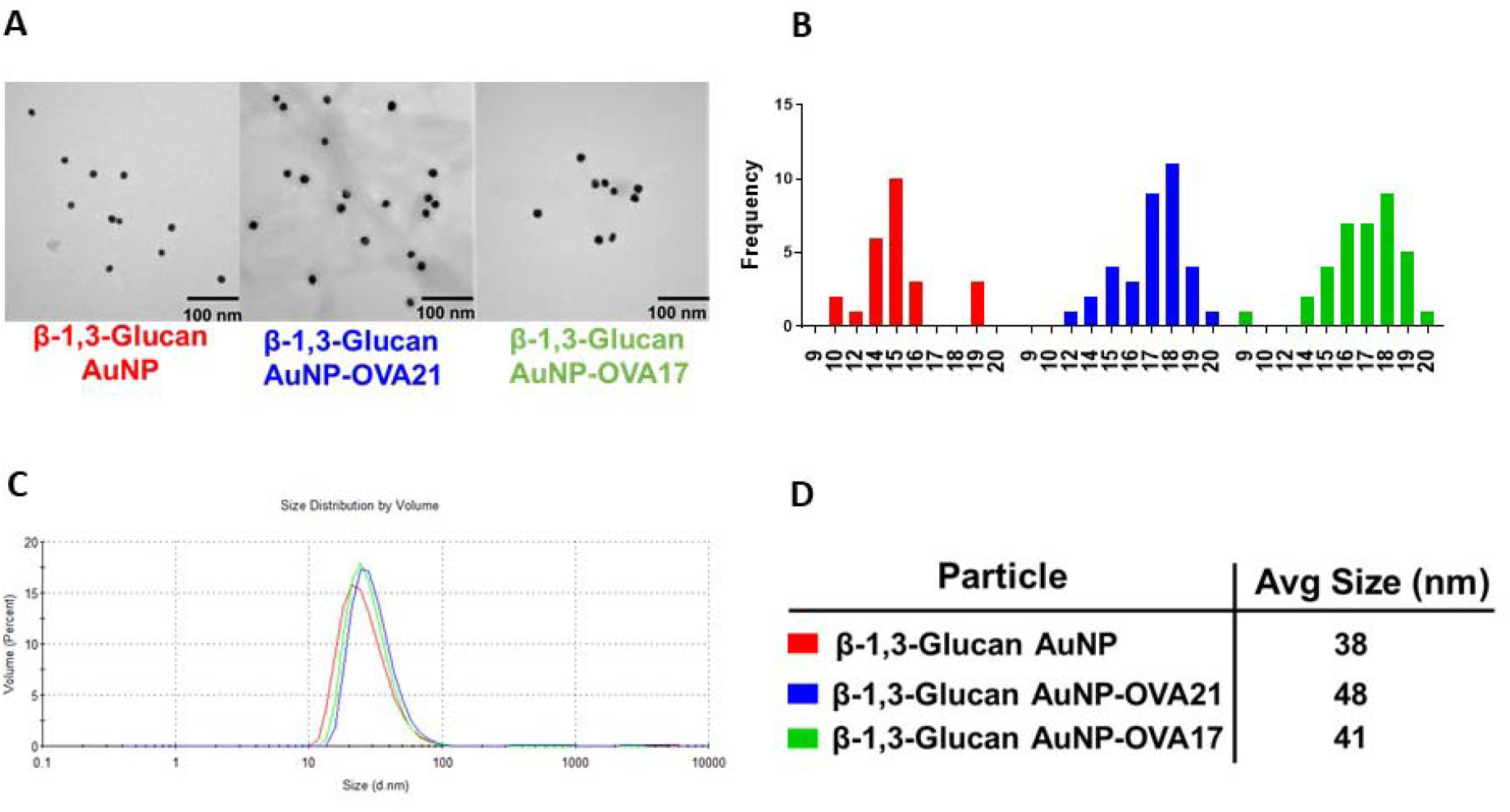
Representative characterization data for B13G-AuNPs and those conjugate with OVA peptides. (A) Transmission electron micrographs; (B) Size histograms for the TEM data in (A). (C) Dynamic Light Scattering volume distribution for B13G-AuNPs from (A); (D) Hydrodynamic diameter of particles from (A) determined from DLS data in (C).

We performed carbohydrate analysis via a simple phenol-sulfuric acid assay to determine the sugars concentration on the particle. Total carbohydrate (measured as glucose monomers) was found to be 2 mmol sugar/AuNP (Figure S5).

### Conjugation of Peptide/Glycopeptide Sequences to B13Gs-Coated AuNPs and Estimation of Peptide Coverage

The attachment of the actual antigens to our newly synthesized B13Gs AuNPs was performed by a simple place exchange reaction with our thiol-containing linked constructs. We were initially unsure if this would be successful considering the probable high coverage of the gold surface by the glycan polymer. However, coating with peptides or glycopeptides went smoothly as we were able to confirm the addition of these antigens by several indirect methods. As described above, for the unconjugated particle, we performed carbohydrate analysis POST-conjugation to observe any displacement of the polymer. We saw an approximate 50% drop in carbohydrate concentration after addition of the antigens, suggesting a displacement of polymer from the gold surface. This is expected as the polymer coating is through non-covalent interaction interactions in contrast to the stronger dative-type bond (~40 kcal/mol)^79^ formed between thiol and gold when antigen is conjugated. However, this “competition” for binding site on the gold surface did not affect the bioactivity of the particles as shown below.

Estimates of the peptide coverage was qualitatively made by the displacement of a fluorescent thiolated peptide that was conjugated to the B13Gs-coated particles. For this we used a commercially available 5kDa FITC-PEG thiol that we conjugated to the B13Gs AuNP. This in turn displaced a portion of the B13Gs from the particle, while the attachment caused the well-known fluorophore quenching by the Forster Resonance Energy Transfer (FRET) properties of 3-dimensional self-assembled monolayers of gold. Subsequent release of the PEG-fluorophore by treatment with dithiothreitol (DTT) restored fluorescence which was quantitated at 525 nm. This corresponded to a loading of the FITC-PEG 362 nM per a solution of 400 μg/mL of nanoparticle (Figure S6).

### Dectin-1 Binding

The B13G AuNPs were compared with free B13Gs for binding to Dectin-1. As shown in Figure S7, ELISA assays where the polysaccharide was bound to the wells and binding was analyzed with biotinylated Dectin-1, B13Gs-loaded particles were shown to bind equally well or better than soluble B13Gs in a dose dependent manner. Binding was observable down to single digit nanomolar concentrations. This result showed that the binding to the targeted C-type lectin was recapitulated in the designed B13Gs-stabilized particles.

### Vaccination studies with B13Gs AuNPs: *in vitro* Model Study with OVA

Before attempting any *in vivo* experiments, initial evaluation of the B13Gs AuNPs with a model system was performed as an *in vitro* pre-screen for appropriate biological activity. Our design took advantage of the availability of a macrophage/T-cell clone pair that can present and recognize a specific Ovalbumin peptide, respectively. Tumor macrophage Dectin-1-expressing cell line P388D1 paired with the Do-11.10 T-cell clone which expresses a T-cell receptor (TCR) that recognizes the 17-mer ovalbumin sequence were chosen for an exploratory assay with our B13Gs AuNPs. In brief, uptake and presentation of the OVA peptide by P388D1 cells within the context of MHC-II will allow recognition by the T-cell clone and release of IL-2. We synthesized this 17-residue peptide encompassing the recognition domain containing OVA amino acids 323-339 (i.e., in single amino acid code: ISQAVHAAHAEINEAGR) and coupled the N-terminus to our thiol-containing linker for conjugation to B13Gs AuNPs (see Figure 3A for a description of the experiment). We also synthesized a second peptide containing a protease recognition domain -GLGR- after the final isoleucine at the N-terminus directly before linker attachment. The peptides were characterized by NMR and both ESI and MALDI mass spec (see Supporting Information for all characterization data). Each of these were coupled to B13G-AuNPs as described above. Figure 3 shows the IL-2 readout resulting from incubation of the OVA-conjugated particles with both the P388D1 and Do-11.10 cells as described in Materials and Methods. As shown in Figure 3B, at 15 μg well, the B13Gs-AuNP-OVA-21 construct (containing the -GLGR- cleavage sequence) was as active as the peptide alone at 200 μg/well (positive control), while the construct without the cathepsin cleavage motif (B13Gs-AuNP-OVA17) was about half as active as the OVA-21 construct. These results suggested that the particles function to enter dectin-1 expressing APCs and retain the ability to present peptide (glycopeptide) cargo to T-cells.

**Figure 3.**
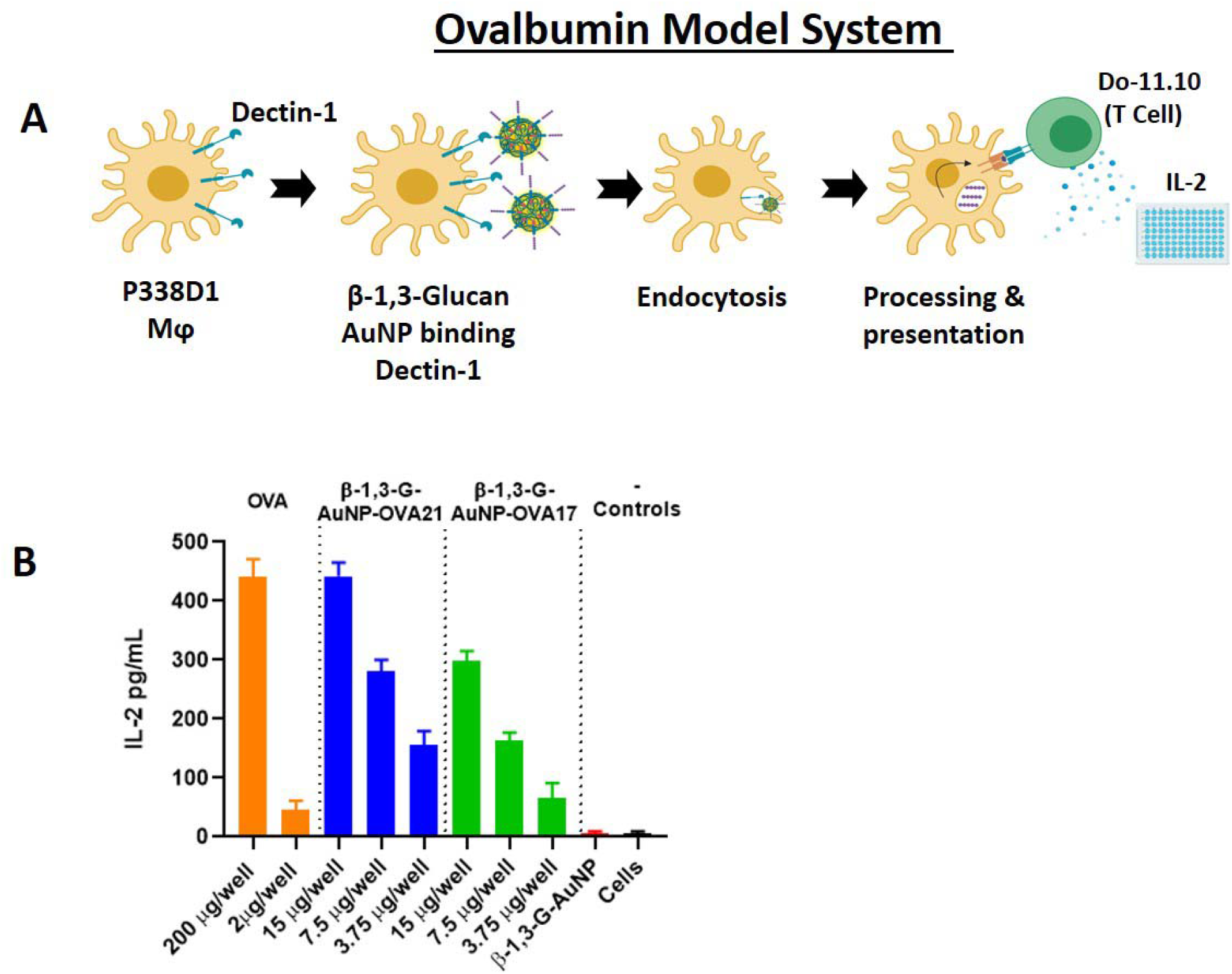
(A) Schematic scheme of the OVA model system. (B) Readout of IL-2 release fom treatment of P338D1 macrophages with various controls and AuNPs, followed by incubation with OVA-specific T-cell clone Do-11.10.

### *in vivo* Vaccination Studies with B13Gs-MUC4/Glycopeptide-Loaded AuNPs

Based on the ovalbumin study, we prepared B13Gs-AuNPs with our MUC4 peptide/glycopeptides from Figure 1. These studies were performed in two stages: First, we prepared B13Gs-AuNPs conjugated with the unglycosylated MUC4 peptide and immunized with two distinct adjuvants to determine the most efficient combination for immune enhancement; Second, the TF-Ser^5^ glycopeptide was conjugated to B13Gs-AuNPs utilizing the adjuvant chosen in the first vaccination. In this step the glycopeptide was also conjugated to the a highly immune-stimulating protein carrier CRM197, a recombinant, non-toxic form of diptheria toxin used as a carrier protein for many polysaccharides^80–82^, as a “positive” control. This was done to compare the new platform to one known to elicit very powerful immune responses to many different haptens. All synthetic haptens were prepared with the -GFLG- tetrapeptide cathepsin cleavage site based on the superior performance of these constructs in the model study. See Figure 4 for a general description of the experimental protocol.

**Figure 4.**
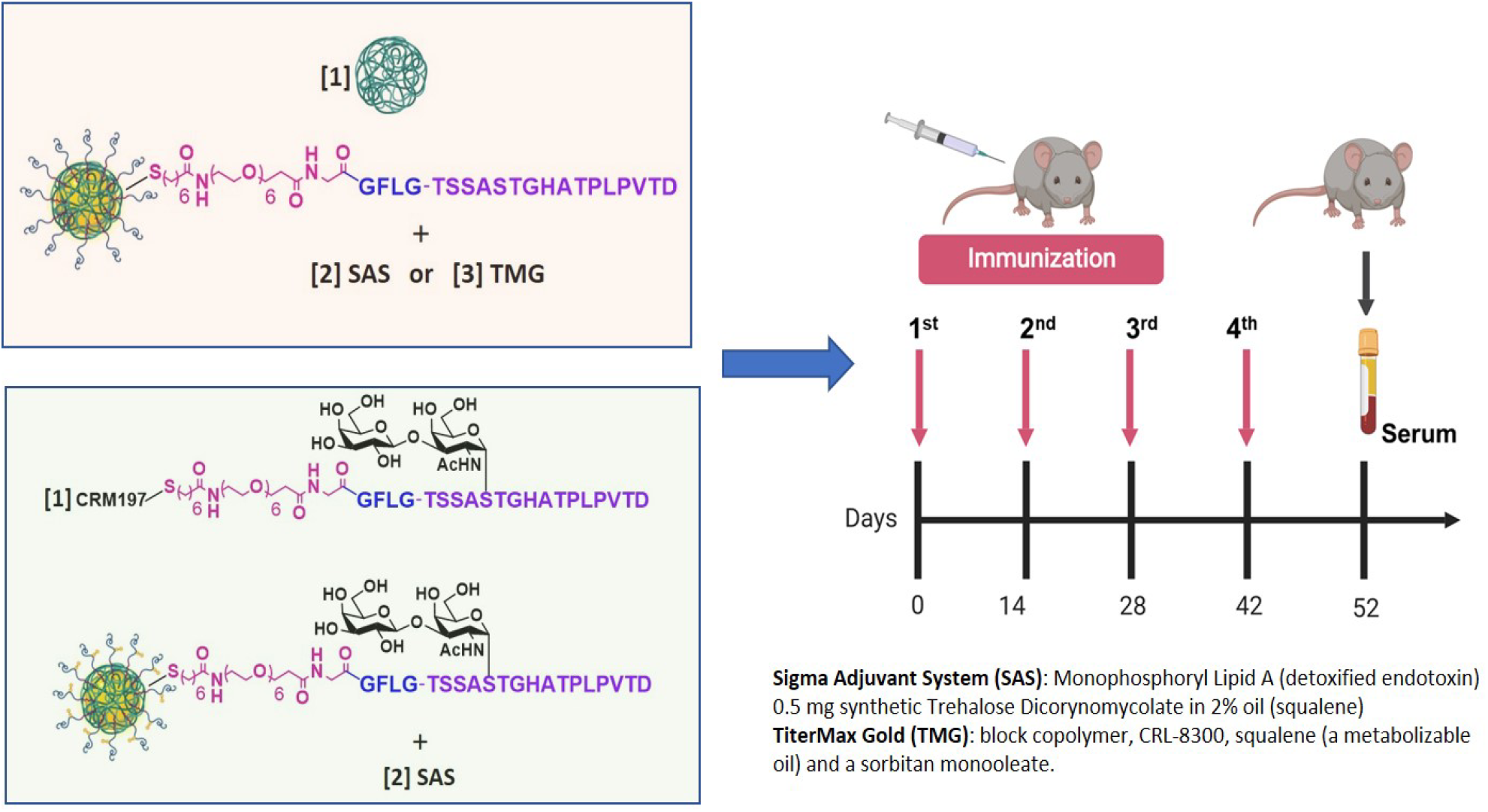
Protocol for in vivo immunizations. In stage 1 (pink box) animals were injected with either B13G-AuNPs or MUC4-B13G-AuNPs with either the Sigma Adjuvant System or TiterMax Gold as adjuvants (descriptions on lower right) with immunization schedule shown on the right, with serum collected at day 52. In stage 2 (light green box), the glycopeptide-coated particles, TF-MUC4-B13G-AuNPs were injected with SAS, along with CRM197-TF-MUC4 conjugate (no nanoparticles). Each stage contained a control group of mice injected with PBS.

Each stage consisted of three groups of 5 mice to be injected. Stage 1 groups were B13G-AuNPs (negative control), MUC4 peptide-conjugated B13G-AuNPs (2 groups, *vide infra*) and PBS. Stage 2 groups were B13Gs-AuNPs (negative control), MUC4 Ser^5^ glycopeptide-conjugated B13Gs-AuNPs and PBS. The experimental design is summarized in Figure 4. Stage 1 used three sets of mice, one PBS control and one set for each of two separate adjuvants: 1) the Sigma adjuvant system, (SAS), which consists of monophosphoryl lipid A (a detoxified endotoxin) and synthetic trehalose dicorynomycolate in a 2% squalene oil base and 2) TiterMax Gold (TMG) which is a mixture of a block copolymer (CRL-8300) and sorbitan monooleate, also in a squalene base. Interestingly, only immunization with SAS elicited both humoral (Figures 5A-C) and cell-based immune response. Both IgG and IgM titers against the MUC4 epitope were generated, with values as high 180,000 for IgG. The IgG subtypes generated were primarily IgG1 and IgG2b isotypes. Control particles gave no response and the TMG vaccination only showed very low IgM titers of any immunoglobulin type.

**Figure 5.**
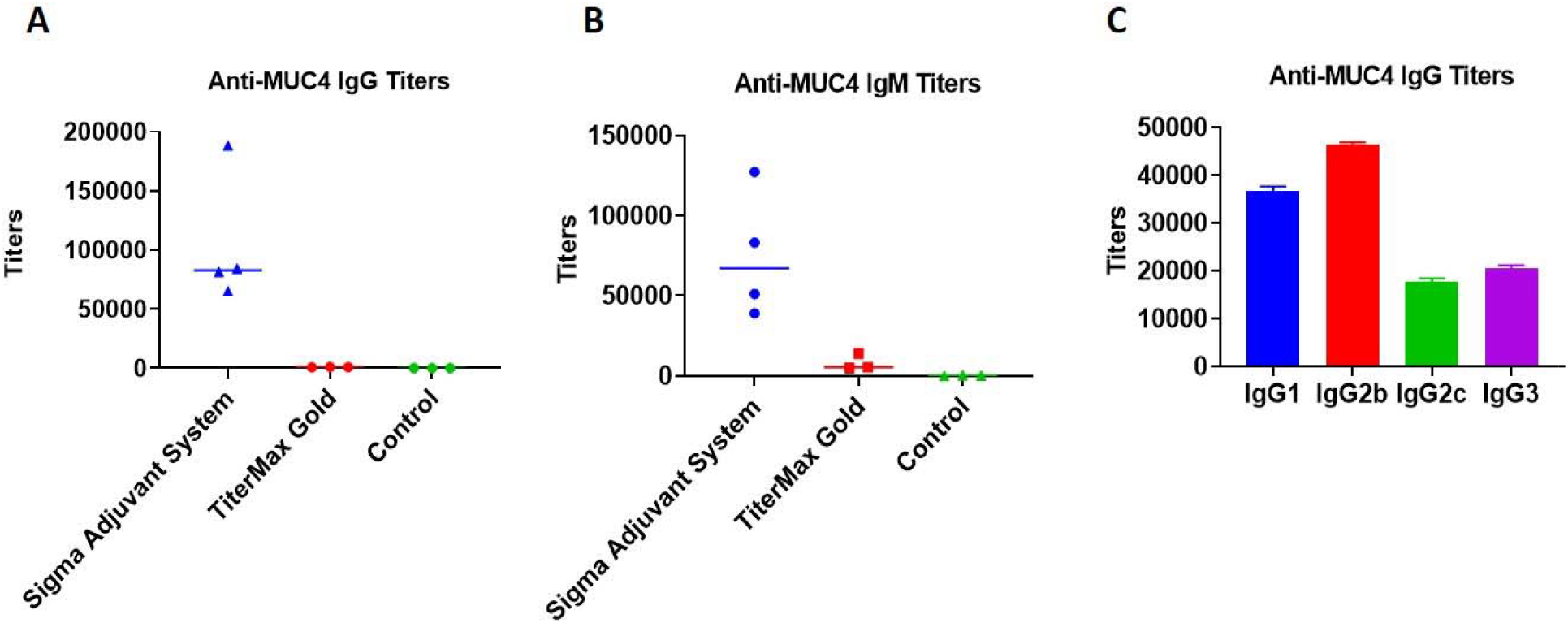
Graph of antibody titers to the MUC4 16-mer peptide from stage 1 immunizations. (A) IgG, (B) IgM and (C) IgG subtypes.

The SAS vaccine subgroup was analyzed by a cytokine bead assay. The repertoire of cytokines expressed can be used as a measure of a cell-mediated response and to stratify the T-helper subtypes that were generated. The SAS adjuvant subset showed stimulation of IL-1β, IL-5, IL-6, Interferon-γ, IL-17, IL-21, IL-23 and MIP3α. Only IL-6 and IL-10 production was seen in the TMG subset from this pilot study, indicating a more immmunosuppressive (IL-10) environment in these mice (See Figures S8-S11). Due to this undesirable outcome and the overall low levels of cytokines expressed in the TMG-adjuvanted mice, it was decided to use only SAS in the follow-up trial with the TF-MUC4 glycopeptide and any other subsequent vaccinations trial with glycopeptide antigens.

The MUC4-TF-Ser^5^-conjugated B13G-AuNP study proceeded identically to the MUC4 peptide study in terms of immunization frequency and amounts of epitope injected. For this study, we uses MUC4-TF-Ser^5^-conjugated B13G-AuNPs, B13G-AuNPs without glycopeptide conjugation and one group vaccinated with a MUC4-TF-Ser^5^-CRM197 conjugate (formed by coupling our thiolated antigen to CRM197-Maleimide, purchased from Fina Biosolutions, LLC; see Figure S12 for a MALDI mass spectrum of the conjugate). All the vaccinations were performed with SAS adjuvant. The CRM197 conjugate generated a very intense immune response with antibody titers as high as 800,000 (Figure S13). Our B13Gs-AuNP conjugates also generated very respectable immunoglobulin responses, comparable to those seen with the unglycosylated peptide nanoparticles (IgG titers as high as 300,000). These results suggested that the platform we are using can generate high humoral immune responses without the need for highly immunogenic carrier protein (See Discussion).

The cell-mediated response was also comparable to the conjugate (1), with cytokine generation of similar quantities as those from peptide-alone vaccinations. While the Th2 response was similar to conjugate 1, the IFN-γ quantities were about 2-fold higher than those seen for conjugate 1. Elispot analysis of splenocytes from sacrificed animals (See Figure S14) showed proliferation of both peptide and glycopeptide-specific T-cell clones, all suggesting that these nanoparticles can be taken up by APCs and their cargo presented T-cells (Figure 6). Important to both vaccination studies was the absence of Il-10 protein expression elicited by either of our vaccines. IL-10 is a cytokine that suppresses the ability of DCs to stimulate CD4^+^ T-cell proliferation and reduces IFN-γ production.^83, 84^ In addition, only the CRM197 conjugate sera contained TGF-β, another cytokine that can have both immunosuppressive and anti-apoptotic effects. Importantly, a trial of the pancreatic cancer vaccine GVAX showed potentiated antitumor activity when combined with TGF-β blockade.^85^

**Figure 6.**
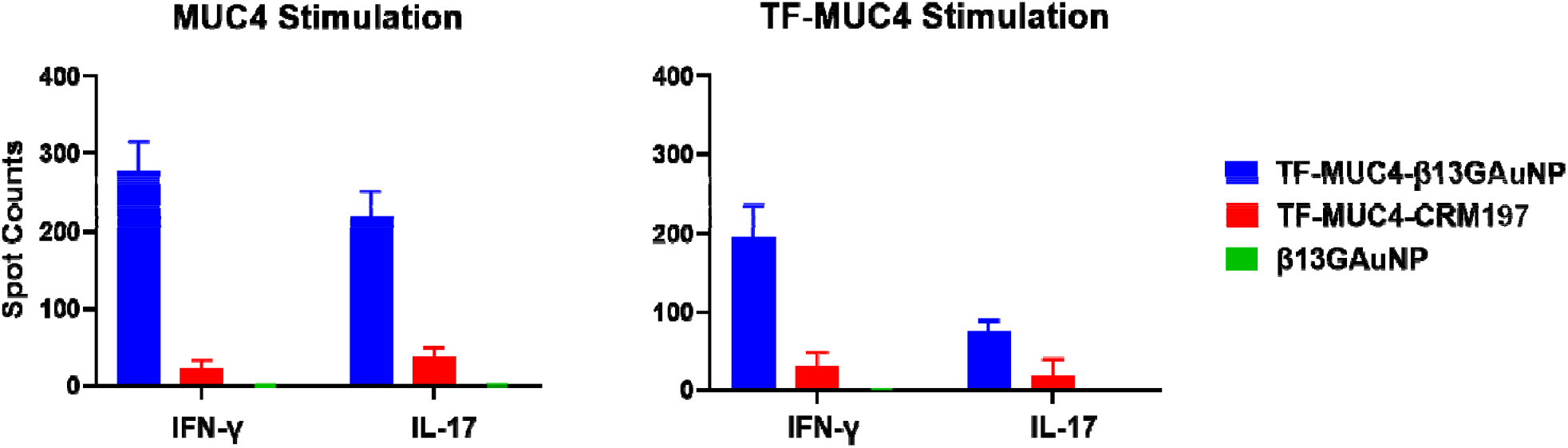
Graphs of ELISspot disk counts comparing vaccinations of our TF-MUC4 B13G AuNP conjugate the TF-MUC4-CRM197 conjugate and control B13G AuNPs when panned for interferon-γ and IL-17 production. While the CRM197 conjugate has an intense humoral response, the TF-MUC4 vaccine construct elicited a much stronger CD4^+^ T-cell response, whereas the control AuNPs consistently showed no response.

## DISCUSSION

In this study we have prepared a AuNP that combines an APC targeting moiety and a novel glycopeptide antigen. This antigen was chosen since we had previously shown that an antibody raised to this glycopeptide was tumor specific.^70^ The use of B13G polymers as both stabilizing and targeting agents has produced a nanoparticle that is adaptable and multifunctional. The microwave-assisted synthesis of the B13G-AuNPs allowed for reproducible production of size-controlled nanoparticles that have a long shelf life, and these constructions may also be functionalized with antigens of various molecular families. Although we did not perform any direct imaging or Dectin-1 knockout studies, both *in vitro* and *in vivo* data showed that these AuNPs can generate distinct and powerful immune responses, strongly suggesting that these particles can be endocytosed by APCs and presented to T-cells to produce mature antibodies and helper T-cells. The preliminary data shown here on the quality, magnitude and uniqueness of the immune response is very encouraging. First, the antibody response was quite respectable for a simple nanoparticle system and compares well with other TACA-based vaccine antibody titers that were generated from either the fully synthetic platforms of Boons, et al.^86, 87^, the Q-beta-based platforms of the Huang lab^34^ or the MUC4 glycopeptide Tetanus Toxoid conjugates of the Kunz group.^88^ The work from Kunz et al., is most relevant to the present work, as this is the only other group that has used MUC4 tandem repeat TACA-containing glycopeptide units in their work. They report titers of 50,000-400,000 for two vaccinations which is very comparable to that observed in the present study; in addition, IgG2 are the predominant isotype of the antibodies produced in the present study. Here we also report results on the cytokine profiles that were generated by presentation of antigen to T-cells by what we assume is an MHC-II-mediated process. Expression of of IL-1β, IL-5, IL-6, Interferon-γ, IL-17, IL-21, IL-23 and MIP3α. were observed in the cytokine array indicating a mixed Th1/Th2/Th17 profile (see Table 1). The expression levels of IL-21 are important because it is known to stimulate levels of NK/NKT and CD8^+^ cells that have proved essential in killing of certain tumors^89–91^ as well as viral-infected cells.^92–94^ The profiles were similar for both the peptide and glycopeptide vaccinations, and the antibody responses were highly specific for our MUC4 constructs, much as the mAb produced by standard KLH-conjugate in a prelude to the present study.^70^

**Table 1.** Secreted cytokine amounts (in picograms) detected in sera of mice vaccinated with our MUC4 glycopeptide B13G gold nanoparticle or a conjugate of that same glycopeptide with CRM197.

|  | Vaccination Epitope |  |  |  |
| --- | --- | --- | --- | --- |
|  | B13G-TF-MUC4-AuNP/SAS | TF-MUC4-CRM | B13G-AuNP | PBS |
| Cytokine Detected | Amount (pg) |  |  |  |
| IL-1β | 140.5 | 1440 | 23.7 | 28.8 |
| IL-5 | 476.75 | 7567.5 | 56.3 | 30.2 |
| IL-6 | 421.5 | 447 | 43 | 25 |
| IL-17 | 107 | 1346 | 42 | 36.2 |
| IL-21 | 103 | 63 | 18.5 | 12.5 |
| IL-23 | 394 | 5811 | 79.8 | 35.3 |
| INF- $\gamma$ | 724 | 7558 | 116.3 | 60.3 |
| MIP-1α | 560 | 5166 | 310 | 215 |
| IL-10 | 0 | 9 | 33.5 | 20.75 |
| TGF-β | 0 | >70,000 | 0 | 0 |

The data from the cytokine array suggests that the CRM-197 conjugates elicit the expression of high circulating levels of cytokines compared to the B13G-AuNP (Table 1). However, from ELISpot analysis, when CD4+ T-cells and DCs were stimulated with MUC4, there was an 8-fold increase in quantifiable spots for IFN-y and IL-17 for B13G conjugates compared to CRM-197. When stimulated with TF-MUC4, there was a reduction in CD4+ activation in comparison to MUC4 stimulation, but the B13G conjugate demonstrated a 4-fold increase of cytokine stimulation. The decrease in CD4+ stimulation towards the glycopeptide can be attributed to possible reduced lysosomal processing of a modified (glycosylated) peptide or lower affinity for the CD4+ T-cell receptor to the antigen-MHCII complex due to “interference’ from the disaccharide. These data bode well for use of simple and widely available materials to construct a vaccine that will raise proper immune responses and whose formulation does not involve the use of detoxified^82^ proteins or large, multiepitope-containing carrier molecules^18^ that can redirect and dampen the immune response to a chosen immunogen.

The synthesis of the B13G-AuNPs is very simple and modular. It is important to note that this procedure only worked well under microwave conditions. This is possibly due to maintenance of solubility of the yeast B13Gs during the reduction or an acceleration of the rate of reaction to facilitate Au^+3^ to Au^0^ reduction and nanoparticle growth seeding.^95^ While it is well known that naked AuNPs as well as citrate stabilized particles are quite sensitive to aggregation/flocculation under various conditions, passivation with appropriate entities can stabilize AuNPs to a variety of conditions, allowing for use in relative “harsh” milieus (i.e., human serum).^96, 97^ The success of any AuNP therapy is contingent on this inherent or instilled stability, and our particles have fit this first and critical criterion. Many reports have shown that polysaccharides can act as reducing/coating agents in the synthesis of AuNPs.^98^ The use of gums such as gellan^69^, karaya^67^ and katira^68^ have been used in what are considered “green” syntheses of AuNPs where reactive reductants and additional stabilizers are unnecessary. There has been one report of the synthesis of both gold and silver nanoparticles using B13Gs as the reductant and passivating agent.^64^ In this study, curdlan, which is a linear (no 1,6 branching) B13G produced as an exopolysaccharide from certain *Rhizobeaceae* species^99^, was used as well as microwave catalysis; however, no biological data was reported and the quality and size of the particles differed from those presented here.

In addition, B13Gs themselves are known to form self-assembled nanoparticles and these have unique applications in the biomedical field.^63, 100, 101^ Some of these particulate forms of B13Gs have also been used to stabilize nanoparticles for macrophage targeting.^63^ In fact, there are many forms of B13Gs from various sources that have been used in research toward immunological enhancement for some time. B13Gs are found primarily in yeast and other fungal species as well as in oats, barley and other cereals and are made up of β-1,3 glucose linkages with various β-1,6 branching points (fungal), whereas the cereal glucans have both β-1,3 and β-1,4 linkages. Curdlan is a linear β-1,3 glucan found in bacteria that has been used in a variety of physical and biochemical studies. Laminarin is a seaweed-derived B13Gs made up of about a 3:1 ratio of 1,3 and 1,6 linkages.^102^ All of these variants have in some form been explored as immune stimulating entities. Targeting through B13Gs has been shown to enhance the vaccination efficacy of polysaccharide antigens in anti-bacterial vaccine design.^48, 103^ In addition, B13Gs can assume a variety of structural forms depending on experimental conditions.^72–74, 76, 77, 104^ Interestingly, B13Gs can adopt a triple helix conformation which is relevant to interaction of this polysaccharide and cellular receptors like Dectin-1.^105^ While not completely characterized in the present work, it can be assumed that the B13Gs on the AuNP surface adopt a conformation that allows proper interaction with Dectin-1 for signal transduction and endocytosis. The processing of the B13Gs before attaching to AuNPs involved dissolution in base and microwave heating to affect reduction and AuNP formation. While this protocol may unravel a tertiary structural element such as a triple helix, reconstitution of a bioactive conformation is assumed to be facilitated by the “bottom-up” synthesis of self-assembled 3-Dimensional gold nanospheres. Structural and conformational studies of the on-particle B13G molecules and the relationship these elements have on activity is currently in progress.

In conclusion, we have prepared a robust and simple platform that can target APCs with various families of antigens for antibody production and T-cell activation in a mouse. The simple conjugation and delivery of *glyco*peptide antigens is highly relevant, as the design of novel therapies based on TACA-containing tandem repeat sequences from tumor-associated mucins is still a very active area of tumor vaccine research; however, no viable products have yet advanced past the various stages of clinical evaluation. The use of a non-toxic, gold nanoparticle platform, combined with pathogen-associated molecular patters that recognize innate PRRs on APCs is an approach that potentially can solve many of the issues associated with vaccine constructs design to date. The ability to deliver relevant antigens through simple peptide and linker chemistry is also an advantage to the current method. The combination of chemoenzymatic TACA synthesis and simple conjugation techniques will allow for the delivery of many different glycopeptide-type antigens with the potential for true immunotherapy against specific cancers. Work toward defining the actual antitumor activity of our platforms against specific pancreatic tumor models is currently underway and will be presented in due course.

## Acknowledgements

We thank James Kelley of the CBL for help with high resolution mass spectra and Kunio Nagashima and the Electron Microscopy Laboratory, Frederick National Laboratories for TEM images. This project has been funded in whole or in part with Federal funds from the National Cancer Institute, National Institutes of Health, under Contract No. HHSN261200800001E. The content of this publication does not necessarily reflect the views or policies of the Department of Health and Human Services, nor does mention of trade names, commercial products, or organizations imply endorsement by the U.S. Government.

## Author Contributions

K.R.T. conceived the project, designed and performed most of the experiments; KAK helped with experiment optimization and performed stability studies; J.J.B. conceived the project, designed and oversaw all experiments and wrote the paper.

## Additional Information

Supporting Information accompanies this text.

## MATERIALS AND METHODS

### Peptide/Glycopepide Synthesis

Peptides sequences were synthesized using a Liberty Prime^™^ automated microwave peptide synthesizer (CEM Corp., Matthews, NC, USA). Figure 1 shows the MUC4 tandem repeat sequence used, and the abbreviation “MUC4” will be used throughout the manuscript to indicate the 16-residue peptide. A Rink amide resin (loading 0.6 mmol/g) was used as the solid support. Standard couplings of amino acids were carried out at 0.125 M in DMF using DIC/OxymaPure^®^ activation and the corresponding Fmoc-protected amino acid (the synthesis method used is optimized by this activator according to Liberty Prime^™^ recommended operation by CEM). Fmoc removal was performed with 40% pyrrolidine in DMF. Coupling of peracetylated TF-serine or TF-threonine glycoamino acids was performed manually at room temperature for 18 h using HOAt, HATU and 2,4,6-trimethyl pyridine.^71^ After glycoamino acid coupling, the resin-bound peptides were added back to the peptide synthesizer and the sequence was completed as per the initial synthesis, except for the addition of DIEA (0.4 equivalents) in a 0.25 M solution of OxymaPure^®^ in DMF. This modification is also recommended by the manufacturer to buffer the acidity of the Oxyma solution to prevent glycoside hydrolysis. The linker shown in Figure 1 was attached to the N-terminus on-resin using our previously described chemistry.^37, 70^ Peptides were cleaved from the resin using TFA under gentle agitation over a period of 2 h at 25°C in the presence of scavengers (TFA/Triisopropyl silane/Water/DOT (92.5:2.5:2.5:2.5) to avoid oxidation. The majority of TFA was removed by a stream of nitrogen and the crude peptides were precipitated by the addition of ice-cold diethyl ether. The precipitates were centrifuged, dried, and purified by High Performance Liquid Chromatography (HPLC) on reverse phase C18 columns using specific gradients of water and acetonitrile (Solvents A and B), each containing 0.1% TFA. Characterization was by high resolution mass spectrometry and high-resolution NMR.

### Synthesis and Characterization of B13Gs AuNPs

β-1,3(1-6) glucan (Yeast Beta Glucan, Megazyme, Bray, Ireland; 3 mg) was suspended in 4.9 mL MilliQ H_2_O contaning 100 μL of 1 M NaOH. This suspension was heated to 90 °C for 30 minutes in a Biotage Initiator microwave reactor to fully dissolve the β-1,3-Glucan. A portion of this B13G stock solution (1.67 mL) was diluted with 3.11 mL H_2_O and 67 μL 1 M NaOH. This mixture was heated to 90 °C for 5 minutes in the microwave reactor. After this initial heating, 100 μL of 10 mM HAuCl_4_ was introduced into the mixture. The mixture was then microwave-heated to 90 °C for 60 minutes to form the β-1,3-Glucan AuNPs. The final concentrations of reagents were: β-1,3-Glucan (1 mg/mL), HAuCl_4_ (0.1 mg/mL), NaOH (20 mM). β-1,3-Glucan AuNPs were isolated by centrifugation on 50 kDa MWCO centricon filters at 4000 rpm for 10 min. The particles were diluted with water repeatedly washed in this manner (7x), further diluted with water and passed through a 0.45 μm filter. The particles were analyzed by UV/Vis (IMPLEN Nanophotometer NP80) DLS (Malvern Zetasizer Nano-ZS) and carbohydrate analysis (*vide supra* and Supporting Information).

The place exchange reaction to add the linker-conjugated peptide and glycopeptide to the particle was performed by dissolving 1 mg of either MUC4-PEG-SH or TF-MUC4-PEG-SH in 1 mL of water and mixing with 3 ml of the freshly prepared B13G-AuNPs in a glass vial. The reaction was placed in a shaking incubator overnight at 45°C. Ultrafiltration of residual small molecules was affected by ultrafiltration using 50K MW-cutoff spin filters. Filtration was repeated 7 times, washing with Ultrapure MilliQ water at each step.

### Transmission Electron Microscopy

For TEM imaging, carbon film-coated 200 mesh copper grids were glow-discharged to improve hydrophilicity. One drop (~3.5 μL) of the sample dissolved in water was applied to a grid for 30 seconds to allow adsorption. The excess solution was removed with filter paper, and the grid was air-dried. Digital micrographs were collected using an Hitachi H-7650 electron microscope equipped with a 2k CCD camera at an operating voltage of 80 kV.

### Dynamic Light Scattering

DLS data were collected on a Malvern Zetasizer model nano ZS in MilliQ water, with samples diluted in 1.5 ml cuvettes. Runs were performed in triplicate at 90 seconds per run.

### ELISA Assays

TF-MUC4-BSA or MUC4-BSA^70^ were coated on Immulon^®^ Microtiter^™^ 4 HBX 96 well-plates at 1 μg/mL in 0.1 M carbonate buffer (pH 9.2) and incubated for 18 h at 4 °C. Plates were washed three times with 200 μL of washing buffer (1 × PBS buffer with 0.05% Tween^®^ 20) and blocked with 200 μL of 3% BSA for 2 h at room temperature. Serum from vaccinated and control mice were initially diluted at 1:1000 and then serially half-log_10_ diluted and added to wells for a final volume of 100 μL in each well The plates were then incubated for 2 h at 37 °C on an orbital shaker. After incubation, the plates were washed three times with 200 μL of washing buffer. Alkaline phosphatase-linked secondary antibodies (anti-IgM and anti-IgG, Jackson ImmunoResearch) were used to detect primary antibodies bound to either TF-MUC4 or MUC4. The secondary anti-IgM antibodies were diluted 1:1000 while the anti-IgG antibodies were diluted 1:5000. A volume of 100 μL of the respective secondary antibody was placed in each well and incubated for 1 h at 37 °C. The plates were washed three times with 200 μL of washing buffer followed by addition of *p*-nitrophenyl phosphate (PNPP, 1 mg mL) in diethanolamine buffer (pH 9.8) at 100 μL per well and further incubated for 30 min. The optical density was read at 405 nm using a BioTek Synergy HTX Multi-Mode Microplate Spectrophotometer. Titers were determined by regression analysis with dilutions plotted against absorbance. The titer cutoff point value was set at 0.3 which was three times the control titer determination for the PBS-innoculated mice anti-sera. Statistical analysis from ELISAs for experimental groups were compared with the controls using the paired *t* test in GraphPad Prism, version 6.

### Ovalbumin Control Assay

P388D1 macrophage cells (2×10^4^ cells/well) were plated and activated by the addition of 10 ng of IFN-γ and 100 ng of LPS per mL of culture for 24 hr. The media was removed and cells were washed carefully to avoid imparting additional stress to the cells. Stimulation of cells with the positive control OVA peptide (17-mer or 21-mer) was performed both at a high (200 nM) and a low concentration (2 nM). The B13G-AuNP-OVA21 and B13G-AuNPs were then used to stimulate the P388D1 macrophage cells at three concentrations (based on an estimated concentration of conjugate on the nanoparticle, see Supporting Information) at 16 nM, 4 nM, and 2 nM. For the unconjugated B13G-AuNP negative control nanoparticle, the cells were stimulated at 16 nM. At 24 hr post-stimulation, the media was removed and 5×10^4^ Do-11.10 T cells were co-cultured with the antigen primed macrophages for for 24 hr. Cell supernatants were examined for IL-2 release using the Mouse IL-2 Quantikine ELISA kit following manufacturer instructions

### Mouse Immunizations

Immunizations were performed on sets of 5 mice per group. Animals were injected on days 0, 14, 28 and 42 by intraperitoneal (IP) injection with 30 ug of nanoparticles (B13G-AuNP, B13G-AuNP-MUC4 or B13G-AuNP-TF-MUC4) in 100 μl of buffer solution with addition of either 100 μl of Sigma Adjuvant System (SAS) or TiterMax Gold Adjuvants or 1X PBS for control. On day 52, blood serum was obtained via cardiac puncture and the spleens were harvested. The spleens were pooled and homogenized from each of the experimental and control groups. Mouse CD4+, CD8+, and DC’s were isolated from each of the groups via MojoSortTM isolation kits. The isolated cells were then used in ELISpot assays.

Similar to the protocol above, vaccinations with a TF-MUC4-CRM197 conjugate were performed in a similar manner only this time using 3 μg of the CRM197 conjugate in 100 μL of 1X PBS (pH 7.4) along with 100 μL of Sigma Adjuvant System (SAS). On day 52 blood sera and spleens were processed as described above.

### Cytokine Array

Mouse Th1/Th2/Th17 Array Q1 (Raybiotech Cat# QAM-TH17-1) was performed using manufacturers protocols. Pooled neat serum from B13Gs-AuNP, B13Gs-AuNP-MUC4 or B13Gs-AuNP-TF-MUC4 and 1X PBS was diluted 2-fold in 1X PBS. These diluted samples (100 μL each) were incubated on the array plates for 1 h at room temperature. The plates were washed 5 times for 5 min at each step and 80 ul of the biotinylated detection antibody cocktail was added and incubated for 1 h. The array plates were further wash 5x for 5 min each. The Cy3 equivalent dye streptavidin-conjugate (80 μL) was added to each well and incubated in the dark for 1 h at room temperature. Plates were read on a Biotek Synergy plate reader.

### ELISpot assays

Dual-Color ELISpot Mouse IFN-γ/IL-17 kits (R&D systems catalog #ELD5007) were used and the assays were performed using the manufacturers protocols. A 10% FBS IMDM media, supplemented with 1% penicillin-streptomycin, 1% GlutaMAX and 1% β-mercaptoethanol was used to culture the isolated splenocytes. CD4+ T-cells (100 μL, 2.3 x105 cells per well) and DC’s (100 μL, 2.0 x105 cells per well) isolated from experimental mice were incubated with either 20 μg of unglycosylated MUC4 peptide or TF-Ser5-MUC4 glycopeptide antigens to stimulate cytokine release. The plates were incubated for 48 hr at 37°C and 5% CO2. Manufacturers protocols were used to develop the plates and were read on a Immunospot Analyzer.

## References

1. Hakomori, S. I., New Directions in Cancer-Therapy Based on Aberrant Expression of Glycosphingolipids - Antiadhesion and Ortho-Signaling Therapy. Cancer Cell-Mon Rev 1991, 3 (12), 461–470.

2. Feizi, T.; Childs, R. A., Carbohydrate Structures of Glycoproteins and Glycolipids as Differentiation Antigens, Tumor-Associated Antigens and Components of Receptor Systems. Trends Biochem Sci 1985, 10 (1), 24–29.

3. Dabelsteen, E., Cell surface carbohydrates as prognostic markers in human carcinomas. J Pathol 1996, 179 (4), 358–369.

4. Jin, K. T.; Lan, H. R.; Chen, X. Y.; Wang, S. B.; Ying, X. J.; Lin, Y.; Mou, X. Z., Recent advances in carbohydrate-based cancer vaccines. Biotechnol Lett 2019, 41 (6-7), 641–650.

5. Wei, M. M.; Wang, Y. S.; Ye, X. S., Carbohydrate-based vaccines for oncotherapy. Med Res Rev 2018, 38 (3), 1003–1026.

6. Feng, D. Y.; Shaikh, A. S.; Wang, F. S., Recent Advance in Tumor-associated Carbohydrate Antigens (TACAs)-based Antitumor Vaccines. Acs Chemical Biology 2016, 11 (4), 850–863.

7. Amon, R.; Reuven, E. M.; Ben-Arye, S. L.; Padler-Karavani, V., Glycans in immune recognition and response. Carbohyd Res 2014, 389, 115–122.

8. Liu, C. C.; Ye, X. S., Carbohydrate-based cancer vaccines: target cancer with sugar bullets. Glycoconjugate J 2012, 29 (5-6), 259–271.

9. Yin, Z. J.; Huang, X. F., Recent Development in Carbohydrate Based Anticancer Vaccines. J Carbohyd Chem 2012, 31 (1-3), 143–186.

10. Guo, Z. W.; Wang, Q. L., Recent development in carbohydrate-based cancer vaccines. Curr Opin Chem Biol 2009, 13 (5-6), 608–617.

11. Franco, A., Glycoconjugates as vaccines for cancer immunotherapy: Clinical trials and future directions. Anti-Cancer Agent Me 2008, 8 (1), 86–91.

12. Xu, Y.; Sette, A.; Sidney, J.; Gendler, S. J.; Franco, A., Tumor-associated carbohydrate antigens: a possible avenue for cancer prevention. Immunol Cell Biol 2005, 83 (4), 440–8.

13. Toyokuni, T.; Singhal, A. K., Synthetic Carbohydrate Vaccines Based on Tumor-Associated Antigens. Chem Soc Rev 1995, 24 (4), 231-+.

14. Yang, Z. G.; Ma, Y. F.; Zhao, H.; Yuan, Y.; Kim, B. Y. S., Nanotechnology platforms for cancer immunotherapy. Wires Nanomed Nanobi 2020, 12 (2).

15. Hu, J.; Wei, P.; Seeberger, P. H.; Yin, J., Mannose-Functionalized Nanoscaffolds for Targeted Delivery in Biomedical Applications. Chem-Asian J 2018, 13 (22), 3448–3459.

16. Yin, Z. J.; Wu, X. J.; Kaczanowska, K.; Sungsuwan, S.; Aragones, M. C.; Pett, C.; Yu, J.; Baniel, C.; Westerlind, U.; Finn, M. G.; Huang, X. F., Antitumor Humoral and T Cell Responses by Mucin-1 Conjugates of Bacteriophage Q beta in Wild-type Mice. Acs Chemical Biology 2018, 13 (6), 1668–1676.

17. Yin, Z. J.; Comellas-Aragones, M.; Chowdhury, S.; Bentley, P.; Kaczanowska, K.; BenMohamed, L.; Gildersleeve, J. C.; Finn, M. G.; Huang, X. F., Boosting Immunity to Small Tumor-Associated Carbohydrates with Bacteriophage Q beta Capsids. Acs Chemical Biology 2013, 8 (6), 1253–1262.

18. Buskas, T.; Thompson, P.; Boons, G. J., Immunotherapy for cancer: synthetic carbohydrate-based vaccines. Chem Commun 2009, (36), 5335–5349.

19. Thompson, P.; Lakshminarayanan, V.; Supekar, N. T.; Bradley, J. M.; Cohen, P. A.; Wolfert, M. A.; Gendler, S. J.; Boons, G. J., Linear synthesis and immunological properties of a fully synthetic vaccine candidate containing a sialylated MUC1 glycopeptide. Chem Commun 2015, 51 (50), 10214–10217.

20. Tagliamonte, M.; Petrizzo, A.; Tornesello, M. L.; Buonaguro, F. M.; Buonaguro, L., Antigen-specific vaccines for cancer treatment. Hum Vacc Immunother 2014, 10 (11), 3332–3346.

21. Pietersz, G. A.; Pouniotis, D. S.; Apostolopoulos, V., Design of Peptide-Based Vaccines for Cancer. Front Med Chem 2010, 5, 127–166.

22. Pietersz, G. A.; Pouniotis, D. S.; Apostolopoulos, V., Design of peptide-based vaccines for cancer. Curr Med Chem 2006, 13 (14), 1591–1607.

23. Ingale, S.; Wolfert, M. A.; Buskas, T.; Boons, G. J., Increasing the Antigenicity of Synthetic Tumor-Associated Carbohydrate Antigens by Targeting Toll-Like Receptors. Chembiochem 2009, 10 (3), 455–463.

24. Zhou, Z. F.; Lin, H.; Li, C.; Wu, Z. M., Recent progress of fully synthetic carbohydrate-based vaccine using TLR agonist as build-in adjuvant. Chinese Chem Lett 2018, 29 (1), 19–26.

25. Nishat, S.; Andreana, P. R., Entirely Carbohydrate-Based Vaccines: An Emerging Field for Specific and Selective Immune Responses. Vaccines-Basel 2016, 4 (2).

26. Andreana, P. R., Zwitterionic polysaccharide (PS A1) as an immune elicitor for vaccine development. Abstr Pap Am Chem S 2009, 237, 748–748.

27. Krishnamachari, Y.; Geary, S. M.; Lemke, C. D.; Salem, A. K., Nanoparticle Delivery Systems in Cancer Vaccines. Pharm Res-Dordr 2011, 28 (2), 215–236.

28. Zhang, Y.; Lin, S. B.; Wang, X. Y.; Zhu, G. Z., Nanovaccines for cancer immunotherapy. Wires Nanomed Nanobi 2019, 11 (5).

29. Sun, Z. Y.; Chen, P. G.; Liu, Y. F.; Zhang, B. D.; Wu, J. J.; Chen, Y. X.; Zhao, Y. F.; Li, Y. M., Multi-component self-assembled anti-tumor nano-vaccines based on MUC1 glycopeptides. Chem Commun 2016, 52 (48), 7572–7575.

30. Teran-Navarro, H.; Calderon-Gonzalez, R.; Salcines-Cuevas, D.; Garcia, I.; Marradi, M.; Freire, J.; Salmon, E.; Portillo-Gonzalez, M.; Frande-Cabanes, E.; Garcia-Castano, A.; Martinez-Callejo, V.; Gomez-Roman, J.; Tobes, R.; Rivera, F.; Yanez-Diaz, S.; Alvarez-Dominguez, C., Pre-clinical development of Listeria-based nanovaccines as immunotherapies for solid tumours: insights from melanoma. Oncoimmunology 2019, 8 (2).

31. Calderon-Gonzalez, R.; Teran-Navarro, H.; Garcia, I.; Marradi, M.; Salcines-Cuevas, D.; Yanez-Diaz, S.; Solis-Angulo, A.; Frande-Cabanes, E.; Farinas, M. C.; Garcia-Castano, A.; Gomez-Roman, J.; Penades, S.; Rivera, F.; Freire, J.; Alvarez-Dominguez, C., Gold glyconanoparticles coupled to listeriolysin O 91-99 peptide serve as adjuvant therapy against melanoma. Nanoscale 2017, 9 (30), 10721–10732.

32. Parry, A. L.; Clemson, N. A.; Ellis, J.; Bernhard, S. S. R.; Davis, B. G.; Cameron, N. R., ‘Multicopy Multivalent’ Glycopolymer-Stabilized Gold Nanoparticles as Potential Synthetic Cancer Vaccines. J Am Chem Soc 2013, 135 (25), 9362–9365.

33. Salahuddin, N.; Elbarbary, A. A.; Salem, M. L.; Elksass, S., Antimicrobial and antitumor activities of 1,2,4-triazoles/polypyrrole chitosan core shell nanoparticles. J Phys Org Chem 2017, 30 (12).

34. Sungsuwan, S.; Wu, X. J.; Huang, X. F., Evaluation of Virus-Like ParticleBased Tumor-Associated Carbohydrate Immunogen in a Mouse Tumor Model. Method Enzymol 2017, 597, 359–376.

35. Yin, Z. J.; Dulaney, S.; Mckay, C. S.; Baniel, C.; Kaczanowska, K.; Ramadan, S.; Finn, M. G.; Huang, X. F., Chemical Synthesis of GM2 Glycans, Bioconjugation with Bacteriophage Q beta, and the Induction of Anticancer Antibodies. Chembiochem 2016, 17 (2), 174–180.

36. Gibadullin, R.; Farnsworth, D. W.; Barchi, J. J.; Gildersleeve, J. C., GaINAc-Tyrosine Is a Ligand of Plant Lectins, Antibodies, and Human and Murine Macrophage Galactose-Type Lectins. Acs Chemical Biology 2017, 12 (8), 2172–2182.

37. Brinas, R. P.; Sundgren, A.; Sahoo, P.; Morey, S.; Rittenhouse-Olson, K.; Wilding, G. E.; Deng, W.; Barchi, J. J., Jr., Design and synthesis of multifunctional gold nanoparticles bearing tumor-associated glycopeptide antigens as potential cancer vaccines. Bioconjug Chem 2012, 23 (8), 1513–23.

38. Svarovsky, S. A.; Szekely, Z.; Barchi, J. J., Synthesis of gold nanoparticles bearing the Thomsen-Friedenreich disaccharide: a new multivalent presentation of an important tumor antigen. Tetrahedron-Asymmetr 2005, 16 (2), 587–598.

39. Biswas, S.; Medina, S. H.; Barchi, J. J., Synthesis and cell-selective antitumor properties of amino acid conjugated tumor-associated carbohydrate antigen-coated gold nanoparticles. Carbohyd Res 2015, 405, 93–101.

40. Glinskii, O. V.; Li, F.; Wilson, L. S.; Barnes, S.; Rittenhouse-Olson, K.; Barchi, J. J.; Pienta, K. J.; Glinsky, V. V., Endothelial integrin alpha 3 beta 1 stabilizes carbohydrate-mediated tumor/endothelial cell adhesion and induces macromolecular signaling complex formation at the endothelial cell membrane. Oncotarget 2014, 5 (5), 1382–1389.

41. Sundgren, A.; Barchi, J. J., Varied presentation of the Thomsen-Friedenreich disaccharide tumor-associated carbohydrate antigen on gold nanoparticles. Carbohyd Res 2008, 343 (10-11), 1594–1604.

42. Gautam, S. K.; Kumar, S.; Cannon, A.; Hall, B.; Bhatia, R.; Nasser, M. W.; Mahapatra, S.; Batra, S. K.; Jain, M., MUC4 mucin-a therapeutic target for pancreatic ductal adenocarcinoma. Expert Opin Ther Targets 2017, 21 (7), 657–669.

43. Kurtenkov, O.; Innos, K.; Sergejev, B.; Klaamas, K., The Thomsen-Friedenreich Antigen-Specific Antibody Signatures in Patients with Breast Cancer. Biomed Res Int 2018.

44. Almogren, A.; Abdullah, J.; Ghapure, K.; Ferguson, K.; Glinsky, V. V.; Rittenhouse-Olson, K., Anti-Thomsen-Friedenreich-Ag (anti-TF-Ag) potential for cancer therapy. Frontiers in Bioscience - Scholar 2012, 4 S (3), 840–863.

45. Janeway, C. A.; Medzhitov, R., Innate immune recognition. Annu Rev Immunol 2002, 20, 197–216.

46. Bauer, S.; Muller, T.; Hamm, S., Pattern Recognition by Toll-Like Receptors. Adv Exp Med Biol 2009, 653, 15–34.

47. Kaisho, T.; Akira, S., Toll-like receptor function and signaling. J Allergy Clin Immun 2006, 117 (5), 979–987.

48. Mayer, S.; Raulf, M. K.; Lepenies, B., C-type lectins: their network and roles in pathogen recognition and immunity. Histochem Cell Biol 2017, 147 (2), 223–237.

49. Penades, S.; de la Fuente, J. M.; Barrientos, A. G.; Clavel, C.; Martinez-Avila, O.; Alcantara, D., Multifunctional glyconanoparticles: Applications in biology and biomedicine. Nato Sci Peace Sec B 2008, 93–101.

50. Gauthier, L.; Chevallet, M.; Bulteau, F.; Thepaut, M.; Delangle, P.; Fieschi, F.; Vives, C.; Texier, I.; Deniaud, A.; Gateau, C., Lectin recognition and hepatocyte endocytosis of GalNAc-decorated nanostructured lipid carriers. J Drug Target 2021, 29 (1), 99–107.

51. Kleski, K. A.; Trabbic, K. R.; Shi, M. C.; Bourgault, J. P.; Andreana, P. R., Enhanced Immune Response Against the Thomsen-Friedenreich Tumor Antigen Using a Bivalent Entirely Carbohydrate Conjugate. Molecules 2020, 25 (6).

52. Napoletano, C.; Zizzari, I. G.; Rughetti, A.; Rahimi, H.; Irimura, T.; Clausen, H.; Wandall, H. H.; Belleudi, F.; Bellati, F.; Pierelli, L.; Frati, L.; Nuti, M., Targeting of macrophage galactose-type C-type lectin (MGL) induces DC signaling and activation. Eur J Immunol 2012, 42 (4), 936–945.

53. Nuti, M.; Zizzari, I.; Napoletano, C.; Rughetti, A.; Rahimi, H.; Antonilli, M.; Bellati, F.; Di Costanzo, F.; Irimura, T.; Wandall, H.; Clausen, H.; Panici, P. B., Macrophage galactose-type C-type lectin receptor for DC targeting of antitumor glycopeptide vaccines. J Clin Oncol 2011, 29 (15).

54. Bundle, D. R.; Paszkiewicz, E.; Elsaidi, H. R. H.; Mandal, S. S.; Sarkar, S., A Three Component Synthetic Vaccine Containing a beta-Mannan T-Cell Peptide Epitope and a beta-Glucan Dendritic Cell Ligand. Molecules 2018, 23 (8).

55. Decout, A.; Silva-Gomes, S.; Drocourt, D.; Blattes, E.; Riviere, M.; Prandi, J.; Larrouy-Maumus, G.; Caminade, A. M.; Hamasur, B.; Kallenius, G.; Kaur, D.; Dobos, K. M.; Lucas, M.; Sutcliffe, I. C.; Besra, G. S.; Appelmelk, B. J.; Gilleron, M.; Jackson, M.; Vercellone, A.; Tiraby, G.; Nigou, J., Deciphering the molecular basis of mycobacteria and lipoglycan recognition by the C-type lectin Dectin-2. Sci Rep-Uk 2018, 8.

56. Xie, J. H.; Guo, L.; Ruan, Y. Y.; Zhu, H. Y.; Wang, L.; Zhou, L.; Yun, X. J.; Gu, J. X., Laminarin-mediated targeting to Dectin-1 enhances antigen-specific immune responses. Biochem Bioph Res Co 2010, 391 (1), 958–962.

57. Backer, R.; van Leeuwen, F.; Kraal, G.; den Haan, J. M. M., CD8(-) dendritic cells preferentially cross-present Saccharomyces cerevisiae antigens. Eur J Immunol 2008, 38 (2), 370–380.

58. Castro, E. D.; Calder, P. C.; Roche, H. M., beta-1,3/1,6-Glucans and Immunity: State of the Art and Future Directions. Mol Nutr Food Res 2021, 65 (1).

59. Nasrollahzadeh, M.; Shafiei, N.; Nezafat, Z.; Bidgoli, N. S. S.; Soleimani, F., Recent progresses in the application of cellulose, starch, alginate, gum, pectin, chitin and chitosan based (nano)catalysts in sustainable and selective oxidation reactions: A review. Carbohyd Polym 2020, 241.

60. Devi, L.; Gupta, R.; Jain, S. K.; Singh, S.; Kesharwani, P., Synthesis, characterization and in vitro assessment of colloidal gold nanoparticles of Gemcitabine with natural polysaccharides for treatment of breast cancer. J Drug Deliv Sci Tec 2020, 56.

61. Padil, V. V. T.; Waclawek, S.; Cernik, M.; Varma, R. S., Tree gum-based renewable materials: Sustainable applications in nanotechnology, biomedical and environmental fields. Biotechnol Adv 2018, 36 (7), 1984–2016.

62. Devendiran, R. M.; Chinnaiyan, S. K.; Yadav, N. K.; Moorthy, G. K.; Ramanathan, G.; Singaravelu, S.; Sivagnanam, U. T.; Perumal, P. T., Green synthesis of folic acid-conjugated gold nanoparticles with pectin as reducing/stabilizing agent for cancer theranostics. Rsc Adv 2016, 6 (35), 29757–29768.

63. Soto, E.; Ostroff, G., Glucan Particles as Carriers of Nanoparticles for Macrophage-Targeted Delivery. Nanomaterials for Biomedicine 2012, 1119, 57–79.

64. El-Naggar, M. E.; Shaheen, T. I.; Fouda, M. M. G.; Hebeish, A. A., Eco-friendly microwave-assisted green and rapid synthesis of well-stabilized gold and core-shell silver-gold nanoparticles. Carbohyd Polym 2016, 136, 1128–1136.

65. Palma, A. S.; Feizi, T.; Zhang, Y. B.; Stoll, M. S.; Lawson, A. M.; Diaz-Rodriguez, E.; Campanero-Rhodes, M. A.; Costa, J.; Gordon, S.; Brown, G. D.; Chai, W. G., Ligands for the beta-glucan receptor, Dectin-1, assigned using “designer” microarrays of oligosaccharide probes (neoglycolipids) generated from glucan polysaccharides. J Biol Chem 2006, 281 (9), 5771–5779.

66. Sze, D. M. Y.; Chan, G. C. F., Effects of Beta-Glucans on Different Immune Cell Populations and Cancers. Adv Bot Res 2012, 62, 179–196.

67. Gangapuram, B. R.; Bandi, R.; Dadigala, R.; Kotu, G. M.; Guttena, V., Facile Green Synthesis of Gold Nanoparticles with Carboxymethyl Gum Karaya, Selective and Sensitive Colorimetric Detection of Copper (II) Ions. J Clust Sci 2017, 28 (5), 2873–2890.

68. Maity, S.; Sen, I. K.; Islam, S. S., Green synthesis of gold nanoparticles using gum polysaccharide of Cochlospermum religiosum (katira gum) and study of catalytic activity. Physica E 2012, 45, 130–134.

69. Dhar, S.; Mali, V.; Bodhankar, S.; Shiras, A.; Prasad, B. L. V.; Pokharkar, V., Biocompatible gellan gum-reduced gold nanoparticles: cellular uptake and subacute oral toxicity studies. J Appl Toxicol 2011, 31 (5), 411–420.

70. Trabbic, K. R.; Whalen, K.; Abarca-Heideman, K.; Xia, L.; Temme, J. S.; Edmondson, E. F.; Gildersleeve, J. C.; Barchi, J. J., A Tumor-Selective Monoclonal Antibody from Immunization with a Tumor-Associated Mucin Glycopeptide. Sci Rep-Uk 2019, 9.

71. Zhang, Y. L.; Muthana, S. M.; Farnsworth, D.; Ludek, O.; Adams, K.; Barchi, J. J.; Gildersleeve, J. C., Enhanced Epimerization of Glycosylated Amino Acids During Solid-Phase Peptide Synthesis. J Am Chem Soc 2012, 134 (14), 6316–6325.

72. Sletmoen, M.; Stokke, B. T., Review: Higher order structure of (1,3)-beta-D-glucans and its influence on their biological activities and complexation abilities. Biopolymers 2008, 89 (4), 310–321.

73. Okobira, T.; Miyoshi, K.; Uezu, K.; Sakurai, K.; Shinkai, S., Molecular dynamics studies of side chain effect on the beta-1,3-D-glucan triple helix in aqueous solution. Biomacromolecules 2008, 9 (3), 783–788.

74. Yadomae, T., Structure and biological activities of fungal beta-1,3-glucans. Yakugaku Zasshi 2000, 120 (5), 413–431.

75. McIntire, T. M.; Brant, D. A., Observations of the (1 -> 3)-beta-D-glucan linear triple helix to macrocycle interconversion using noncontact atomic force microscopy. J Am Chem Soc 1998, 120 (28), 6909–6919.

76. Bohn, J. A.; BeMiller, J. N., (1->3)-beta-D-glucans as biological response modifiers: A review of structure-functional activity relationships. Carbohyd Polym 1995, 28 (1), 3–14.

77. Saito, H.; Tabeta, R.; Yoshioka, Y.; Hara, C.; Kiho, T.; Ukai, S., A High-Resolution Solid-State C-13 Nmr-Study of the Secondary Structure of Branched (1-]3)-Beta-D-Glucans from Fungi - Evidence of 2 Kinds of Conformers, Curdlan-Type Single-Helix and Laminaran-Type Triple-Helix Forms, as Manifested from the Conformation-Dependent C-13 Chemical-Shifts. B Chem Soc Jpn 1987, 60 (12), 4267–4272.

78. Pooja, D.; Panyaram, S.; Kulhari, H.; Reddy, B.; Rachamalla, S. S.; Sistla, R., Natural polysaccharide functionalized gold nanoparticles as biocompatible drug delivery carrier. Int J Biol Macromol 2015, 80, 48–56.

79. Burgi, T., Properties of the gold-sulphur interface: from self-assembled monolayers to clusters. Nanoscale 2015, 7 (38), 15553–15567.

80. MacCalman, T. E.; Phillips-Jones, M. K.; Harding, S. E., Glycoconjugate vaccines: some observations on carrier and production methods. Biotechnol Genet Eng Rev 2019, 35 (2), 93–125.

81. Dagan, R.; Poolman, J.; Siegrist, C. A., Glycoconjugate vaccines and immune interference: A review. Vaccine 2010, 28 (34), 5513–23.

82. Hsieh, C. L., Characterization of saccharide-CRM197 conjugate vaccines. Dev Biol (Basel) 2000, 103, 93–104.

83. Mittal, S. K.; Cho, K. J.; Ishido, S.; Roche, P. A., Interleukin 10 (IL-10)-mediated Immunosuppression: MARCH-I INDUCTION REGULATES ANTIGEN PRESENTATION BY MACROPHAGES BUT NOT DENDRITIC CELLS. J Biol Chem 2015, 290 (45), 27158–27167.

84. Mittal, S. K.; Roche, P. A., Suppression of antigen presentation by IL-10. Curr Opin Immunol 2015, 34, 22–27.

85. Soares, K. C.; Rucki, A. A.; Kim, V.; Foley, K.; Solt, S.; Wolfgang, C. L.; Jaffee, E. M.; Zheng, L., TGF-beta blockade depletes T regulatory cells from metastatic pancreatic tumors in a vaccine dependent manner. Oncotarget 2015, 6 (40), 43005–43015.

86. Ingale, S.; AWolfert, M.; Gaekwad, J.; Buskas, T.; Boons, G. J., Robust immune responses elicited by a fully synthetic three-component vaccine. Nat Chem Biol 2007, 3 (10), 663–667.

87. Buskas, T.; Ingale, S.; Boons, G. J., Towards a fully synthetic carbohydrate-based anticancer vaccine: Synthesis and immunological evaluation of a lipidated glycopeptide containing the tumor-associated Tn antigen. Angew Chem Int Edit 2005, 44 (37), 5985–5988.

88. Cai, H.; Palitzsch, B.; Hartmann, S.; Stergiou, N.; Kunz, H.; Schmitt, E.; Westerlind, U., Antibody induction directed against the tumor-associated MUC4 glycoprotein. Chembiochem 2015, 16 (6), 959–67.

89. Davis, M. R.; Zhu, Z. W.; Hansen, D. M.; Bai, Q.; Fang, Y. J., The role of IL-21 in immunity and cancer. Cancer Lett 2015, 358 (2), 107–114.

90. Thedrez, A.; Harly, C.; Morice, A.; Salot, S.; Bonneville, M.; Scotet, E., IL-21-Mediated Potentiation of Antitumor Cytolytic and Proinflammatory Responses of Human V gamma 9V delta 2 T Cells for Adoptive Immunotherapy. J Immunol 2009, 182 (6), 3423–3431.

91. Davis, I. D.; Skak, K.; Hunder, N.; Smyth, M. J.; Sivakumar, P. V., Interleukin-21 and Cancer Therapy. Targeted Cancer Immune Therapy 2009, 43–59.

92. Mendez-Lagares, G.; Lu, D.; Merriam, D.; Baker, C. A.; Villinger, F.; Van Rompay, K. K. A.; McCune, J. M.; Hartigan-O’Connor, D. J., IL-21 Therapy Controls Immune Activation and Maintains Antiviral CD8(+) T Cell Responses in Acute Simian Immunodeficiency Virus Infection. Aids Res Hum Retrov 2017, 33, S81–S92.

93. Pallikkuth, S.; Parmigiani, A.; Pahwa, S., The role of interleukin-21 in HIV infection. Cytokine Growth F R 2012, 23 (4-5), 173–180.

94. Sarra, M.; Pallone, F.; Macdonald, T. T.; Monteleone, G., Targeting Interleukin-21 in Immune-Mediated Pathologies. Curr Drug Targets 2010, 11 (5), 645–649.

95. Li-Ping, Y.; Wei-Xia, T., Studies on Au colloidal nanoparticles synthesized by microwave irradiation. Acta Phys-Chim Sin 2006, 22 (4), 513–516.

96. Mukhopadhyay, A.; Grabinski, C.; Afrooz, A. R. M. N.; Saleh, N. B.; Hussain, S., Effect of Gold Nanosphere Surface Chemistry on Protein Adsorption and Cell Uptake In Vitro. Appl Biochem Biotech 2012, 167 (2), 327–337.

97. Daniel, M. C.; Astruc, D., Gold nanoparticles: Assembly, supramolecular chemistry, quantum-size-related properties, and applications toward biology, catalysis, and nanotechnology. Chem Rev 2004, 104 (1), 293–346.

98. Escarcega-Gonzalez, C. E.; Garza-Cervantes, J. A.; Vazquez-Rodriguez, A.; Morones-Ramirez, J. R., Bacterial Exopolysaccharides as Reducing and/or Stabilizing Agents during Synthesis of Metal Nanoparticles with Biomedical Applications. Int J Polym Sci 2018, 2018.

99. Cai, Z. X.; Zhang, H. B., Recent progress on curdlan provided by functionalization strategies. Food Hydrocolloid 2017, 68, 128–135.

100. Liu, Q. Y.; Duan, B. C.; Xu, X. J.; Zhang, L. N., Progress in rigid polysaccharide-based nanocomposites with therapeutic functions. J Mater Chem B 2017, 5 (29), 5690–5713.

101. Sakurai, K.; Uezu, K.; Numata, M.; Hasegawa, T.; Li, C.; Kaneko, K.; Shinkai, S., beta-1,3-glucan polysaccharides as novel one-dimensional hosts for DNA/RNA, conjugated polymers and nanoparticles. Chem Commun 2005, (35), 4383–4398.

102. Okazaki, M.; Adachi, Y.; Ohno, N.; Yadomae, T., Structure-Activity Relationship of (1-]3)-Beta-D-Glucans in the Induction of Cytokine Production from Macrophages, in-Vitro. Biol Pharm Bull 1995, 18 (10), 1320–1327.

103. Qiao, W. L.; Ji, S. Y.; Zhao, Y. B.; Hu, T., Conjugation of beta-glucan markedly increase the immunogencity of meningococcal group Y polysaccharide conjugate vaccine. Vaccine 2015, 33 (17), 2066–2072.

104. Yoshioka, Y.; Uehara, N.; Saito, H., Conformation-Dependent Change in Antitumor-Activity of Linear and Branched (1-]3)-Beta-D-Glucans on the Basis of Conformational Elucidation by C-13 Nuclear-Magnetic-Resonance Spectroscopy. Chem Pharm Bull 1992, 40 (5), 1221–1226.

105. Hanashima, S.; Ikeda, A.; Tanaka, H.; Adachi, Y.; Ohno, N.; Takahashi, T.; Yamaguchi, Y., NMR study of short beta(1-3)-glucans provides insights into the structure and interaction with Dectin-1. Glycoconjugate J 2014, 31 (3), 199–207.

