## Supporting Information for "A Stable Nano-Vaccine for the Targeted Delivery of Tumor-Associated Glycopeptide Antigens"

**Select Experimental Methods**..………………………………………….Page S2-S3

**B3G AuNP Synthesis Reproducibility**………………………………….Page S4

**B13G AuNP Salt and Serum Stability.**…………….……………………Page S4-S5

**TEM Images**…………………………………………………………………Page S5

**Quantitation of B13G Coating**………….………………………………..Page S6

**Quantitation of FITC-Thiol Coating**…....………………………………..Page S6

**Dectin-1 Binding**……………………………………………………………Page S7

**Select Cytokine Profiles**…………………………………………………..Page S7-S8

**CRM197 Conjugate Mass Spectra**…..…………………………………..Page S9

**Antibody Titers**……………………………………………………………..Page S10

**ELISpot Figures**…………………………………………………………….Page S11

**NMR Spectra, Analytical HPLC, Hi Res MS**…………………………...Page S12-S17

**General Experimental** **Procedures**: Routine chemicals were purchased from Sigma-Aldrich. Tetrachloroauric acid (HAuCl4) was purchase from Wuhan Golden Wing Industry & Trade Co, Wuhan China. All amino acids and peptide synthesis material were purchased from CEM corp, (Matthews, NC). Peracetylated TF-Serine glycoamino acid was either prepared as previously described or purchased from Sussex Research, Ottawa, Ontario, Canada. Solvents were dried in a Grubb still percolation system under a nitrogen atmosphere. MALDI mass spectra were collected on a Shimadzu Axima Confidence MALDI-TOF mass spectrometer equipped a high mass CovalX HM4 detector operated in linear positive ion mode. Samples were prepared for MALDI analysis by desalting using a 0.5 mL 30K Amicon Ultra centrifugal filter. Samples were spotted on an Axima 384 well sample plate using the overlayer method with sinapinic acid as the matrix. Dynamic light scattering and zeta potential data were collected on a Malvern Nano-ZS Zetasizer instrument. Proton and carbon NMR data were collected on either a Bruker NanoBay 400 Mhz spectrometer with a Bruker 2-channel SMART probe or on a Bruker AVANCE III 500 MHz spectrometer with a TCI (^1^H, ^13^C, ^15^N) cryoprobe at 25°C. Most data were run in 90%/10% H_2_O/D_2_O. Water suppression was performed using excitation sculpting (1D pulse sequence zgesgp).

**Total Carbohydrate Determination of B13G AuNPs**. For all standards and samples, 50 uL of material was plated on a clear 96 well plate and 150 uL of concentrated H_2_SO_4_ was added. The plate was placed in a 37° C shaking incubator for 30 minutes. Then, 30 uL of a 5% phenol solution was added to each well and the plate was incubated for 30 minutes at 37 C. Plate were read using an OD of 490 nm.

The standard glucose curve was prepared using a two-fold dilution over 7 points ranging from 100-1.56 ug/mL of glucose. For determination of peptide-coated AuNPs, 50 ug/mL of similarly treated B13G AuNP, B13G-OVA21-AuNP and  B13G-OVA17-AuNPs were interpolated off of the standard glucose curve.

**FITC-PEG-SH 5K loading on B13G-AuNP**. One mL of 3 mg/ml solution of FITC-PEG-SH was added to 3 mL of a B13G-AuNP OD=1.0. The reaction was placed in a shaking incubator overnight at 45°C. The solution was concentrated in a spin filter 50K MWC 7 times. A standard curve of FITC-PEG-SH was generated from 100 ul of 0.5 mg solution that was serially diluted. The FITC-B13G-AuNP was examined on a fluorescent intensity plate reader @488 nm. The initial fluorescent intensity was low (due to quenching) and the wells were treated with a solution of DTT (50 mg/mL) and 10 uL was added to the well to displace FITC from gold and fluorescence was read at 488 nm.

**MALDI Mass Spectrometry**. MALDI mass spectra were collected on a Shimadzu Axima Confidence MALDI-TOF mass spectrometer equipped a high mass CovalX HM4 detector operated in linear positive ion mode. The samples were prepared for MALDI analysis by desalting using a 0.5 mL 30K Amicon Ultra centrifugal filter. Samples were spotted on an Axima 384 well sample plate using the overlayer method with sinapinic acid as the matrix. MALDI data for the CRM197 conjugates is shown in Figure S14.

**High Resolution Mass Spectrometry**. LC/MS analysis was carried out in positive ion mode on an Orbitrap LTQ-XL system that was configured with a UV-visible DAD detector in-line. A 2.0-µl aliquot of the analysis solution (1 mg/ml) was injected onto a 2.1 X 50 mm Phenomenex 2.6 µm XB-C18 Kinetex column for separation using LC/MS-grade CH_3_CN and H_2_O in a combination isocratic and linear gradient program, as indicated below, at a flow rate of 250 µl /min. With a rapid solvent reset and equilibration, the complete analysis cycle time was 18 min.

Solvent A – 2% CH_3_CN/H_2_O with 0.1% HCOOH

Solvent B – 90% CH_3_CN/H_2_O with 0.1% HCOOH

0 – 2 min	isocratic at 100% A

2 – 11 min	linear gradient from 0% - 100% B

11 – 14 min	isocratic at 100% B (90% CH_3_CN/H_2_O)

14 – 16 min	linear reset to 100% A (2% CH_3_CN/H_2_O)

16 – 18 min	equilibrate at 100% A

**Dectin-1 Binding**. Clear bottom Immulon 4HBX plates were coated with 50 μL of 50 μg/mL solution of B13G-AuNPs, B13G-AuNPs-MUC4, B13G-AuNP-TF-MUC4 or B13Gs. Plates were incubated at 37 C for 4 h to allow for evaporation. Plates were washed with 1X PBS to remove any unbound material. Fc-hDectin-1 (Invivogen, San Diego, Ca; 100 μL) in concentrations of 100 nM, 33 nM, 11 nM, 3.7 nM, and 1.2 nM was added in triplicate to the aforementioned plate coatings and incubated for 2 hr at 37°C. Plates were washed 3 times with washing buffer. Goat Anti-Human IgG, (Fc fragment specific) HRP conjugate was diluted to 1:5000, (100 μL) of this solution was added to the wells and incubated for 1 h at 37°C. Plates were washed with washing buffer 3 times and 100 uL of TMB liquid substrate system for ELISA was added to each well. Plates were incubated for 10 minutes and the reaction was stopped by adding 50 uL of H_2_SO_4_. Plates were read at 450 nm.

**Analytical HPLC.** Analytical HPLC chromatograms were collected using an Agilent 1100 series analytical HPLC system with an Agilent Eclipse plus C18 (3.5 μm, 4.6X100 mm) column. Lyophilized peptides were reconstituted in LC-MS grade water to 5 mg/mL. 5 μL of sample was injected into the system at a draw speed of 100 μL/min and a flow rate of 1.0 mL/min. A linear gradient was used over the course of 30 minutes beginning at a ratio of 100% water:0% acetonitrile to 20% water:80% acetonitrile.

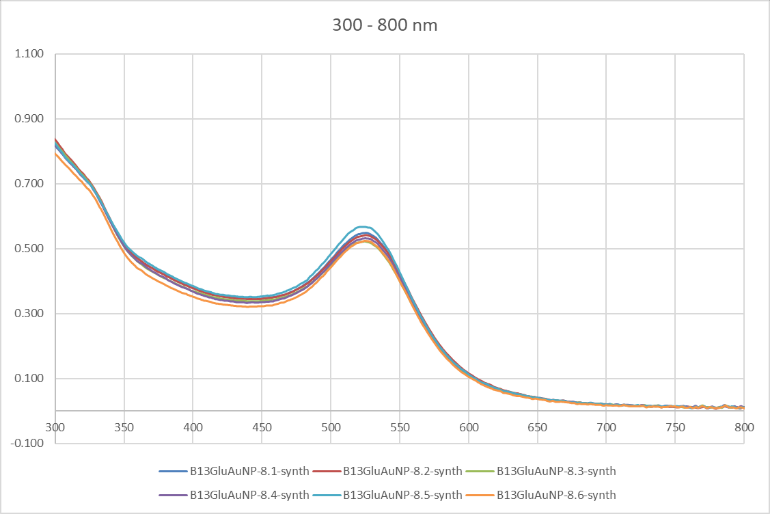

**Z-average and PDI**

8.1 – 38.72 nm (0.208)

8.2 – 36.86 nm (0.220)

8.3 – 38.56 nm (0.242)

8.4 – 37.67 nm (0.213)

8.5 – 38.47 nm (0.198)

8.6 – 41.63 nm (0.212)

**Figure S1.** UV/SPR spectra of six different syntheses (labeled 8.1 – 8.6) of B13G AuNPs. The table to the right depicts the Z-average sizes and polydispersity indices of the six synthetic AuNPs.

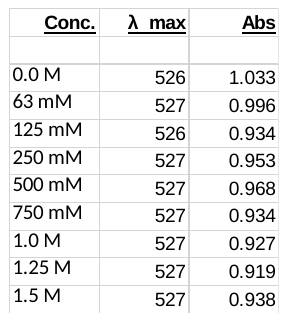

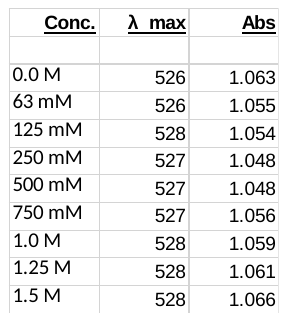

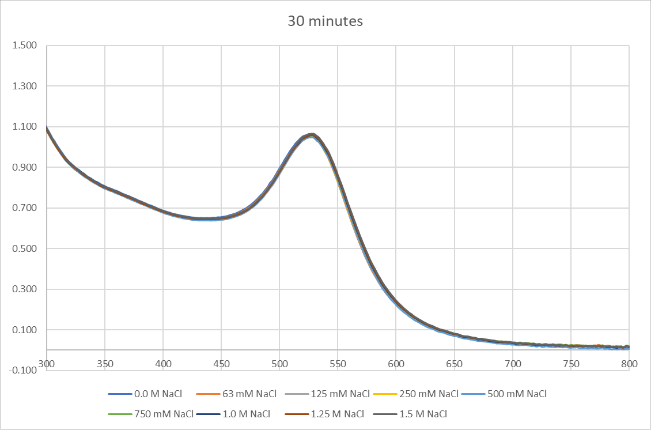

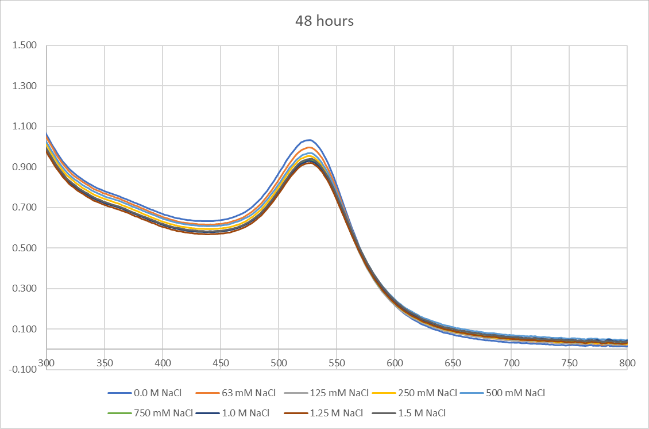

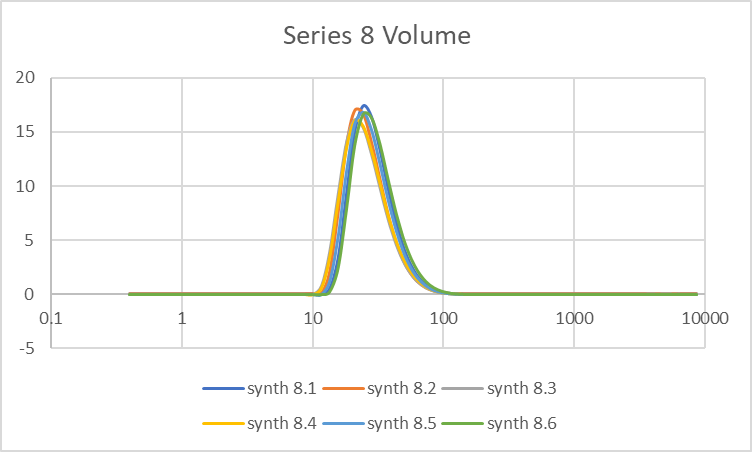

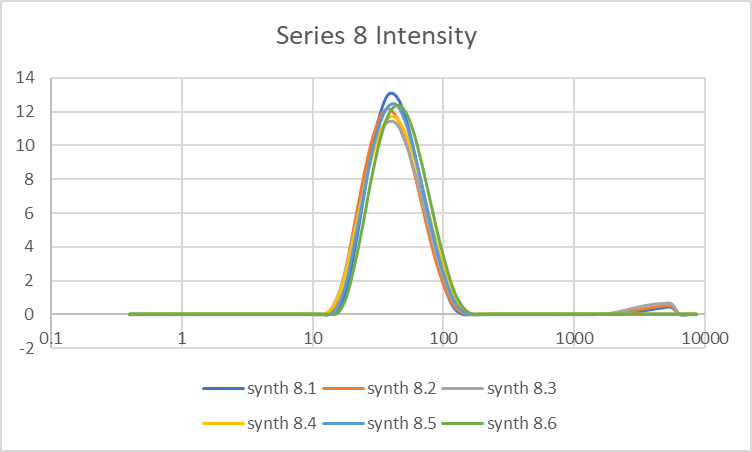

**Figure S2.** UV/SPR and DLS spectra of the same six different syntheses of B13G AuNPs. Top figures show an overlay of UV spectra of B13G AuNPs incubated with increasing concentrations of NaCl (0 – 1.5 M) after 30 min (upper left) and 48 hours (upper right). The tables show the salt concentration, absorbance maxima and raw absorbance units for each experiment. Lower figures: Overlays of DLS intensity data (lower left) and volume data (lower right) from the six synthetic B13G AuNPs.

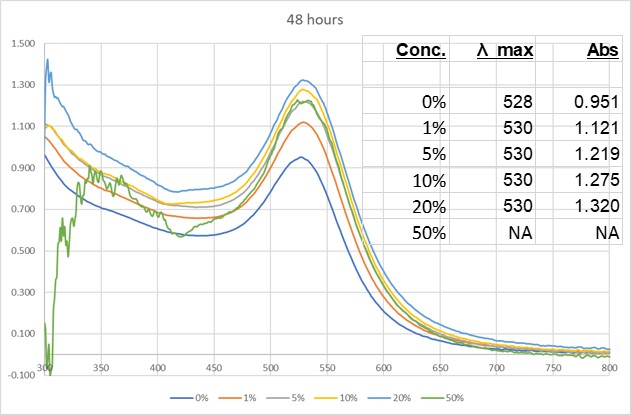

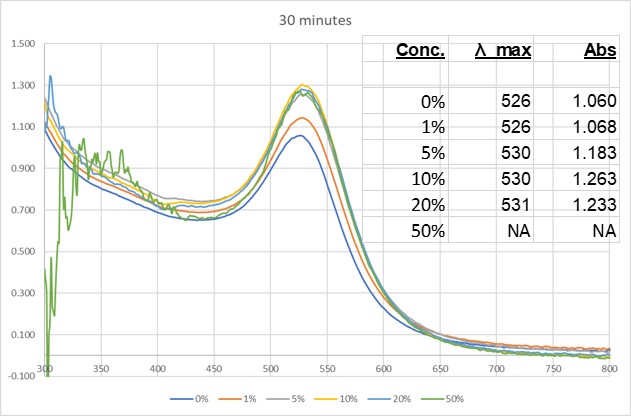

**Figure S3.** UV/SPR of a select synthesis (8.4) of B13G AuNPs after treatment with increasing percentages of human serum (0% - 50%, by volume). The figure on the left shows an overlay of UV spectra after 30 min and the one on the right is after 48 h. The noise in the spectrum of highest percentage serum is due to the increasing turbidity of the solution as serum is increased and not due to any particle aggregation.

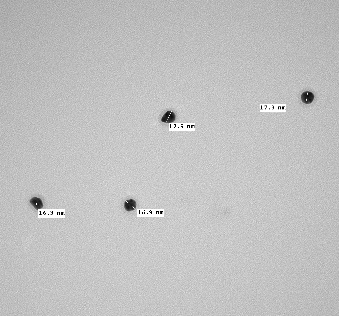

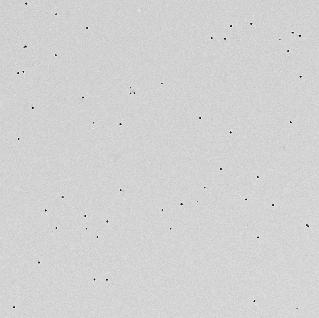

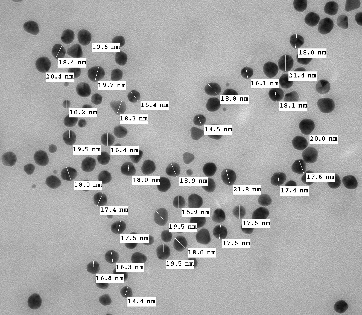
**A	B	C	D**

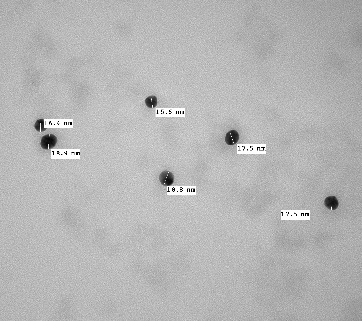

**Figure S4.** Representative TEM images of B13G-AuNP-MUC4 (A, B) and B13G-AuNP-TF-MUC4 (C, D) with size measurements. The sample in **C** is more dilute but particles are very uniform. AuNPs in D are an expansion of the upper right portion of C

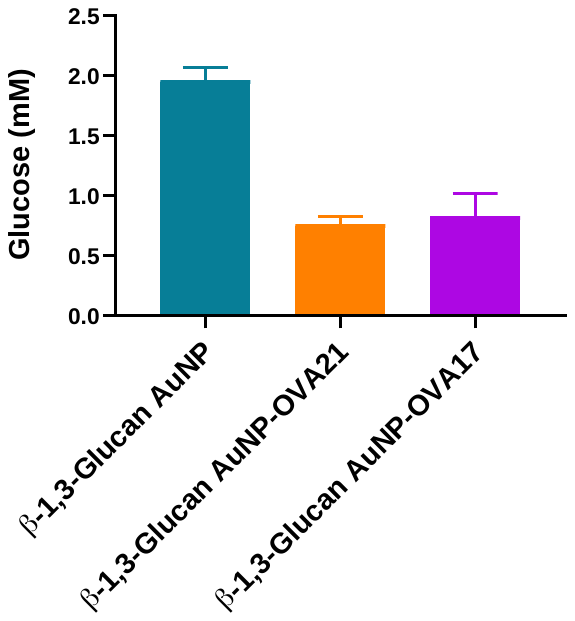

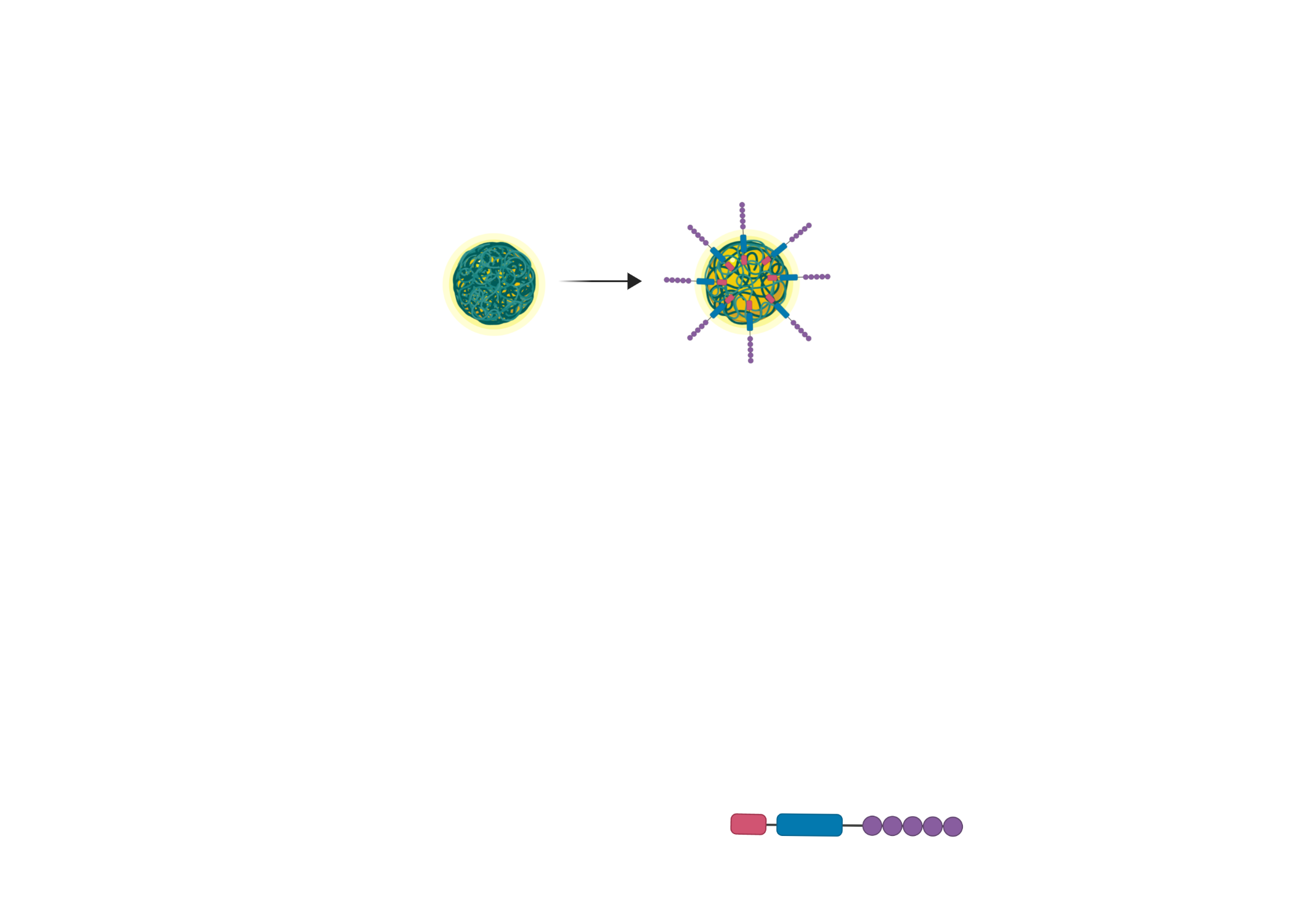

**2 mM**

**.7 mM**

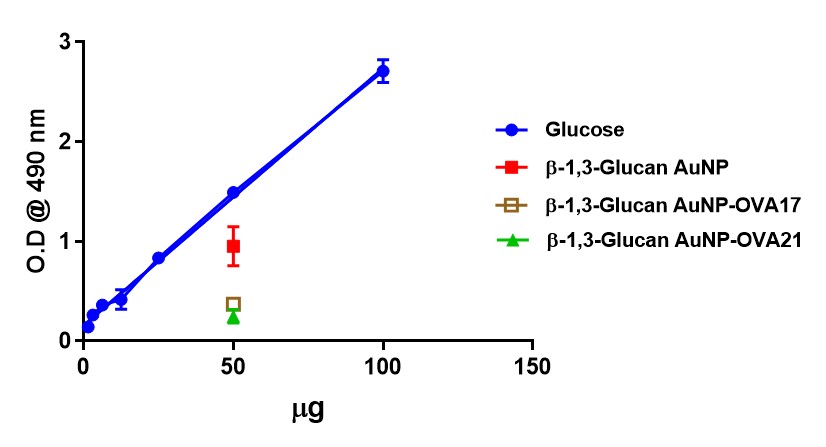

**Figure S5**. Displacement of B13G from particle after place exchange with OVA peptides. Concentration of sugar from the glycopolymer goes from 2 mM to 0.7 mM upon coating. Standard curve of glucose concentration is also shown below

.

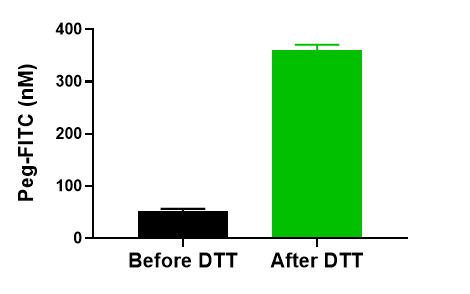

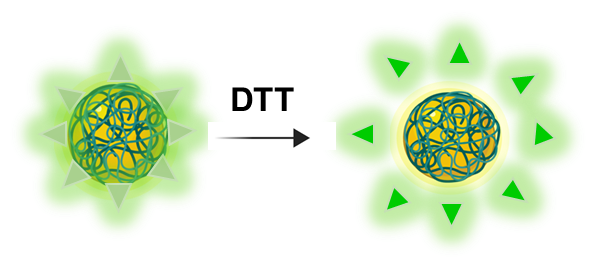

**Quenched**

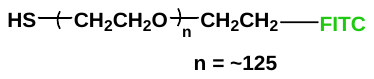

**Figure S6.** Graph of concentration of PEG-FITC-Thiol (structure on lower right) before and after Dithiothreitol treatment. Coupling to AuNPs will quench fluorescence; displacement restores fluorescence and signal is enhanced. Structure of FITC Thiol is

shown in lower right.

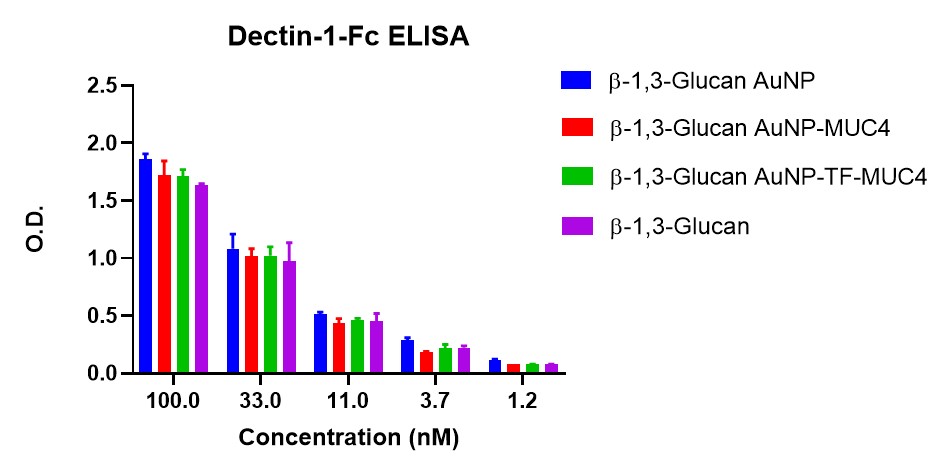
**Figure S7.** Dectin-1 ELISA assay showing binding to various AuNPs used in this study. The C-terminal domain of human Dectin-1 was used conjugated to a human IgG Fc region through a 10 amino acid linker. This allowed for better detection with a Fc-specific human anti-IgG antibody. Dectin1 binding remains even after subsequent coating with antigen peptide/glycopeptide and is detectable down to single digit nanomolar concentrations.

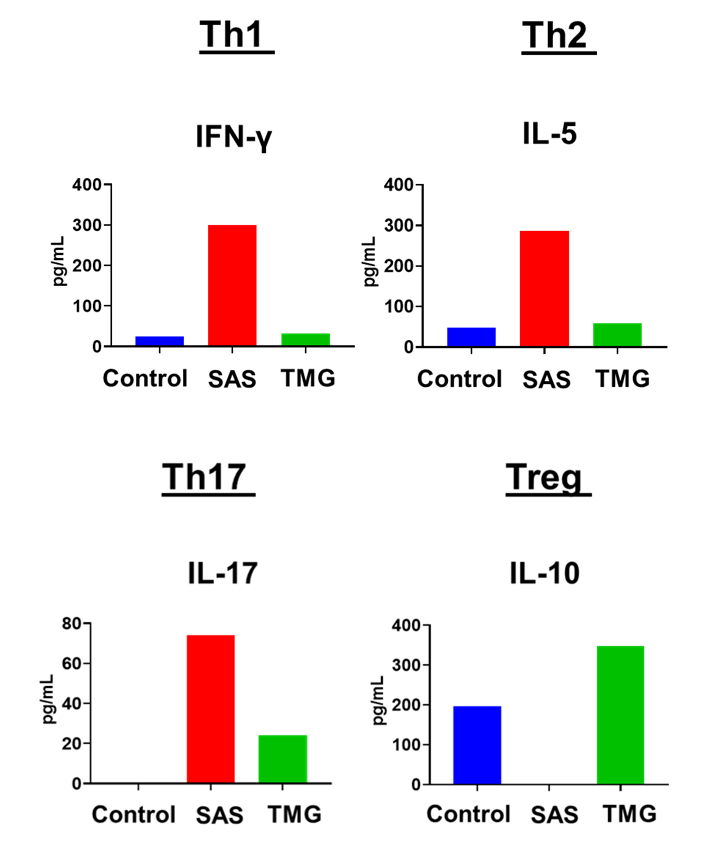

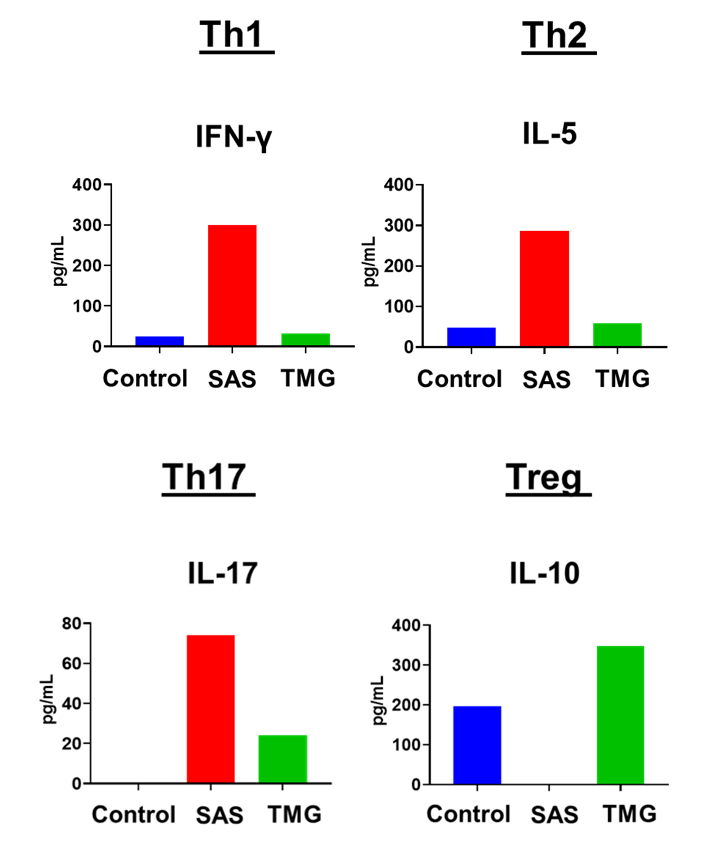

**Figure S8.** Expression of IFN-γ, IL-5, IL-17 and IL-10 after vaccination with MUC4-B13G-AuNP, with a comparison between amounts (pg/ml) generated using SAS or TMG adjuvants. Controls are uncoated B13G AuNPs.

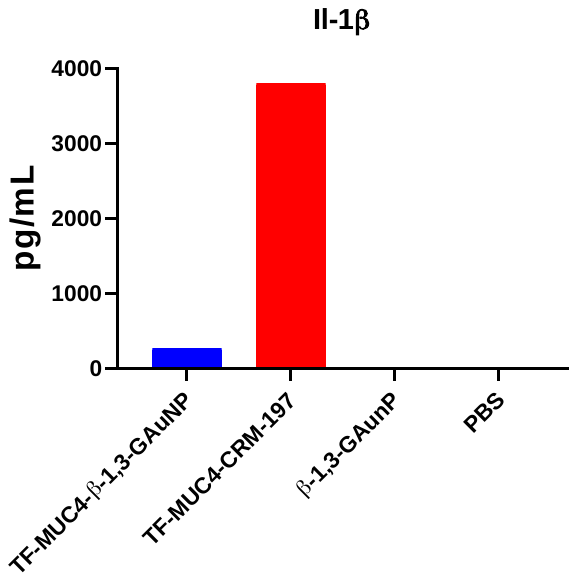

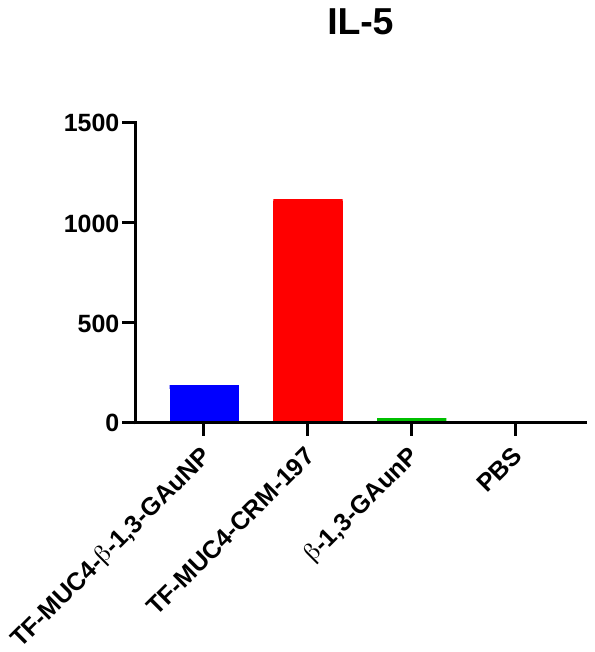

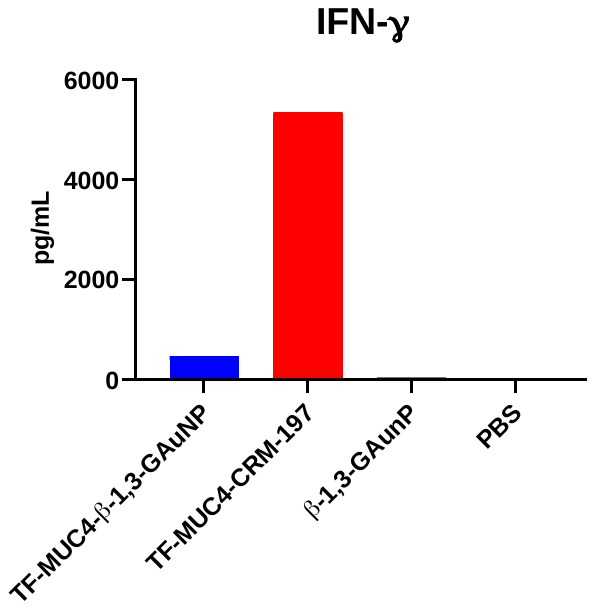

**Figure S9.** Comparison between amounts (pg/ml) of IFN-γ, IL-1β and IL-5 after vaccination with TF-MUC4-B13G-AuNP and TF-MUC4-CRM197 conjugates, both with the SAS adjuvant. Controls are unconjugated B13G AuNPs and PBS.

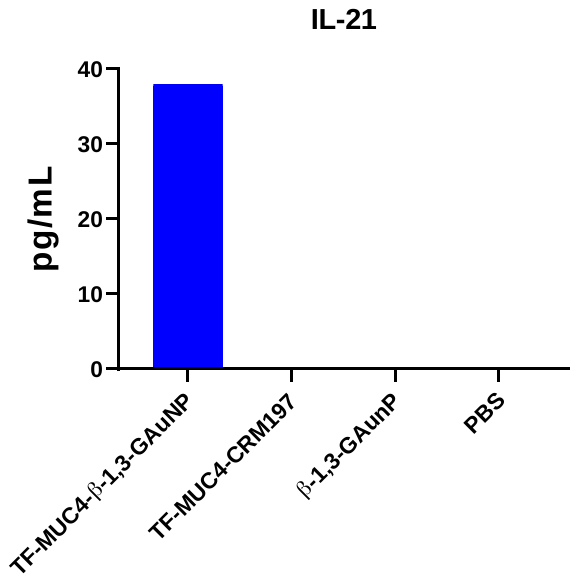

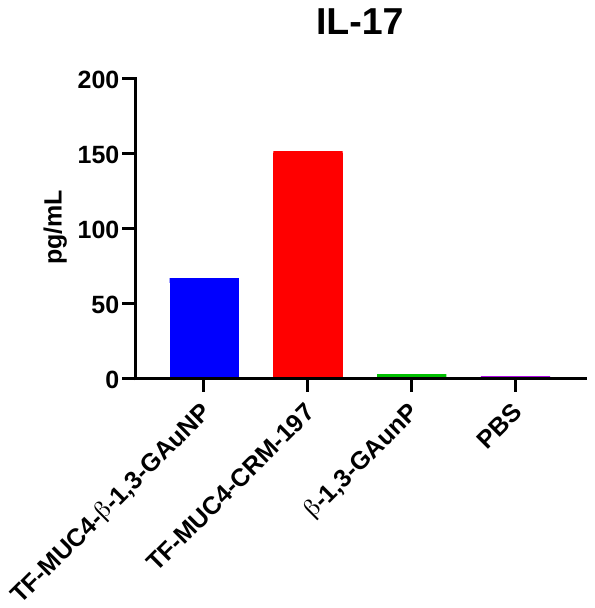

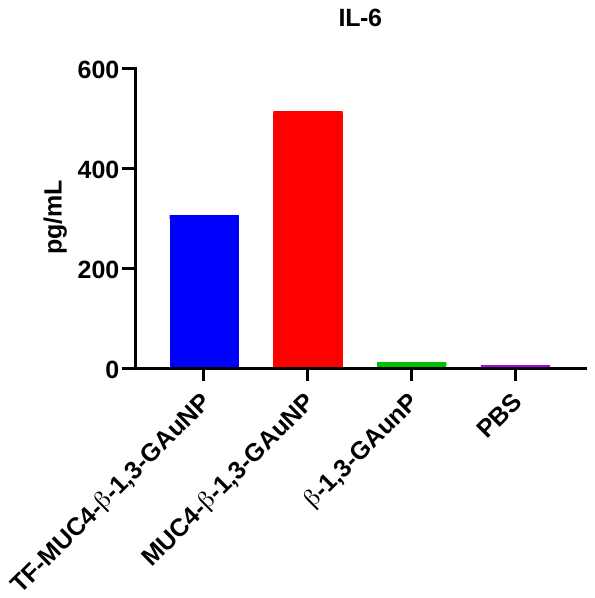

**Figure S10.** Comparison between amounts (pg/ml) of IL-6, IL-17 and IL-21 after vaccination with TF-MUC4-B13G-AuNP and TF-MUC4-CRM197 conjugates, both with the SAS adjuvant. Controls are unconjugated B13G AuNPs and PBS.

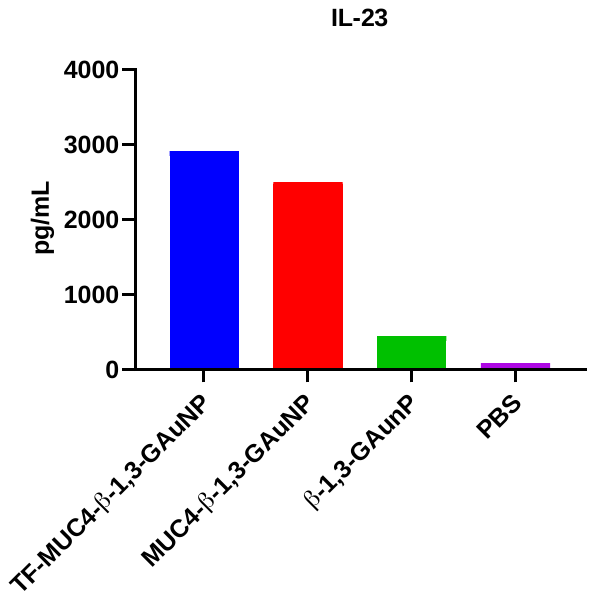

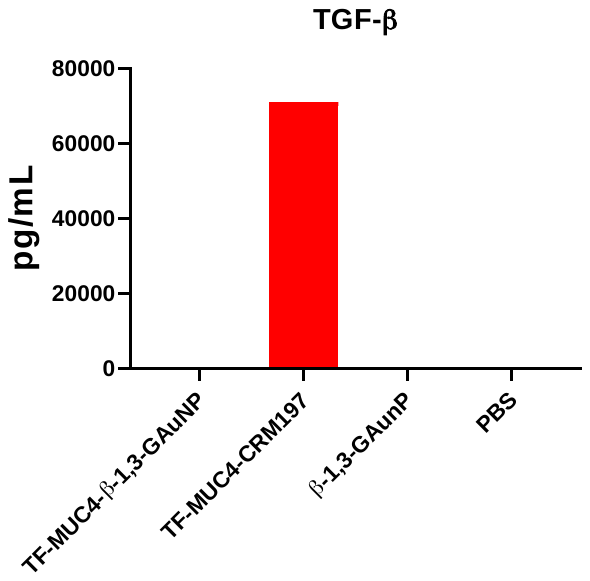

**Figure S11.** Comparison between amounts (pg/ml) of IL-23 and TGF-β after vaccination with TF-MUC4-B13G-AuNP and TF-MUC4-CRM197 conjugates, both with the SAS adjuvant. Controls are unconjugated B13G AuNPs and PBS.

**^A^**

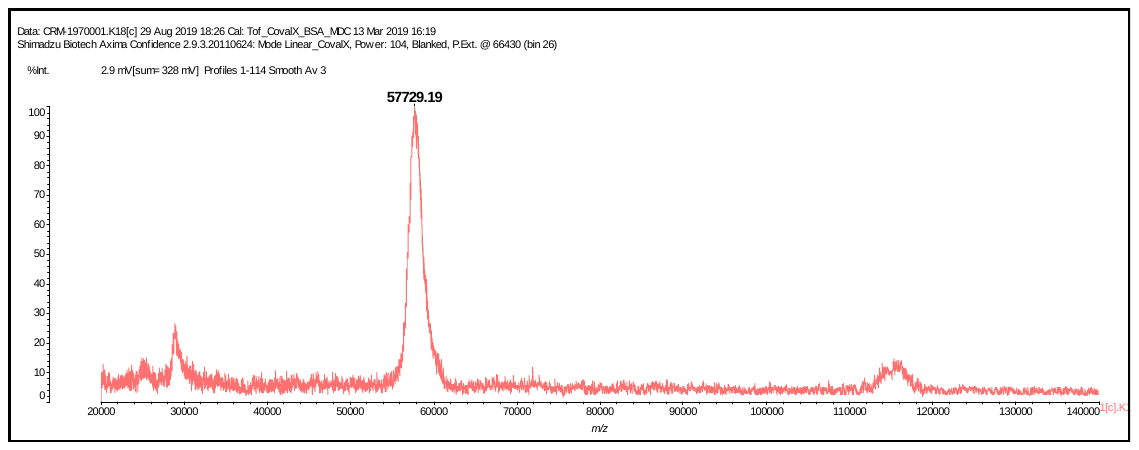

**^B^**

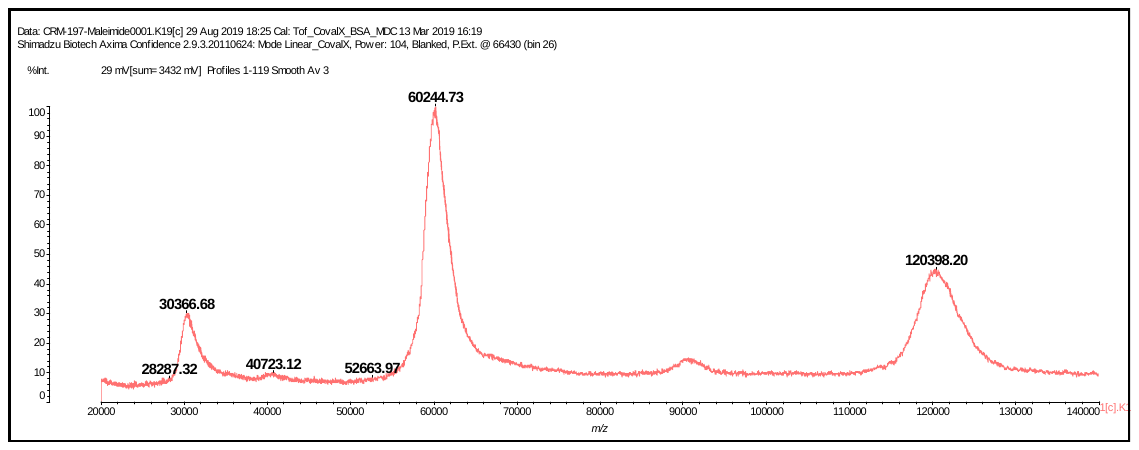

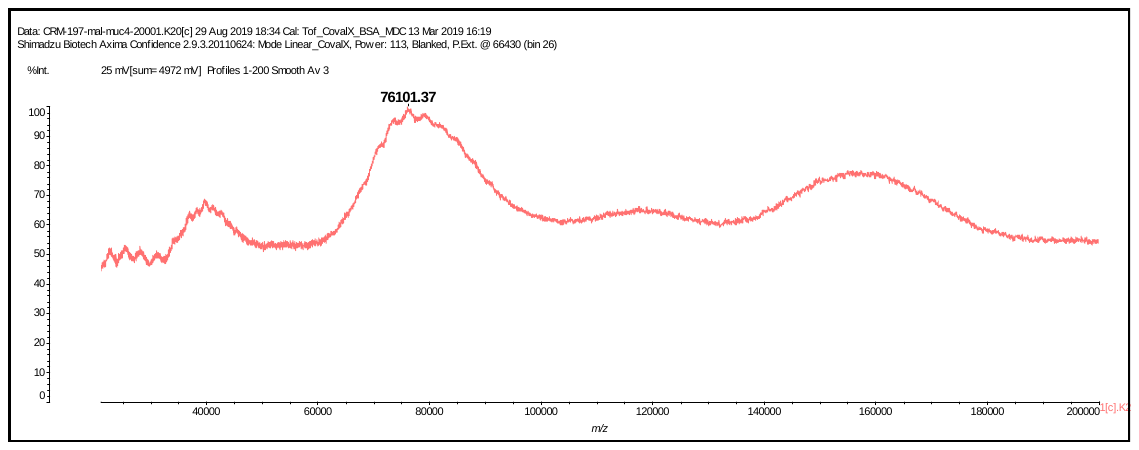
**^C^**

**Figure S12.** MALDI Mass spec spectra of (A) CRM197, (B) CRM197-Maleimide and (C) CRM197-Maleimide-TF-MUC4 conjugate, indicating approximately 8-10 glycopeptides were conjugated to the protein.

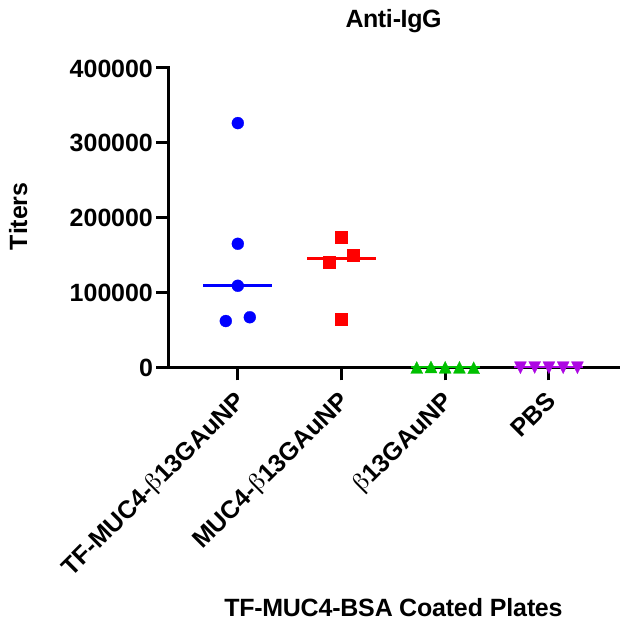

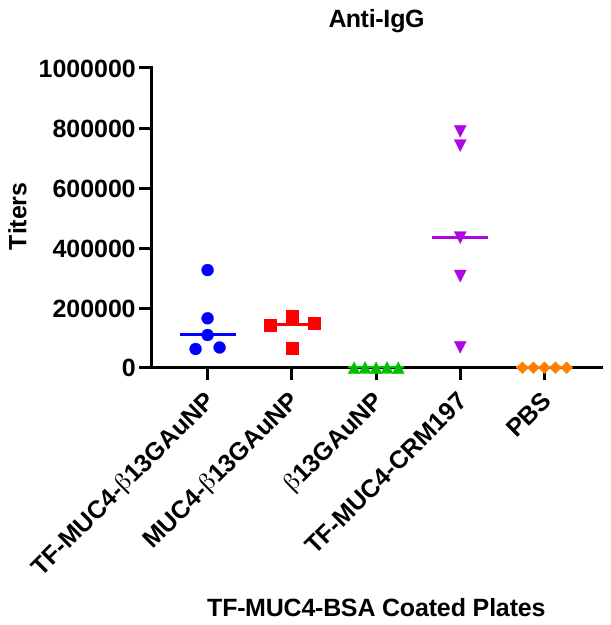

**Figure S13.** IgG Antibody titers generated to vaccinations with glycopeptide-coated B13G AuNPs and to the same glycopeptide conjugated to CRM197. Graph on the right is an expansion of lanes 1, 2, 3 and 5 from the graph on the left

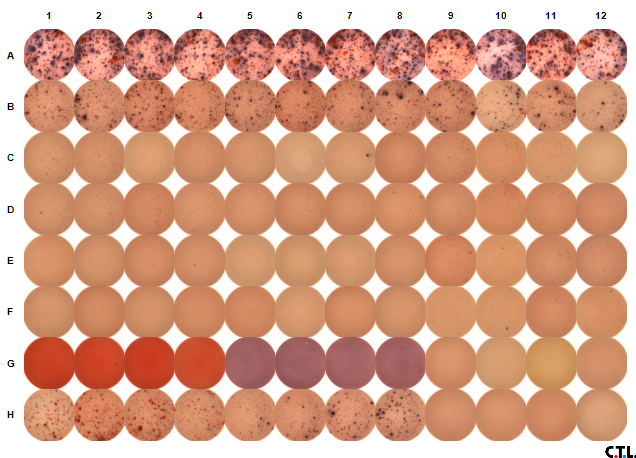

**A**

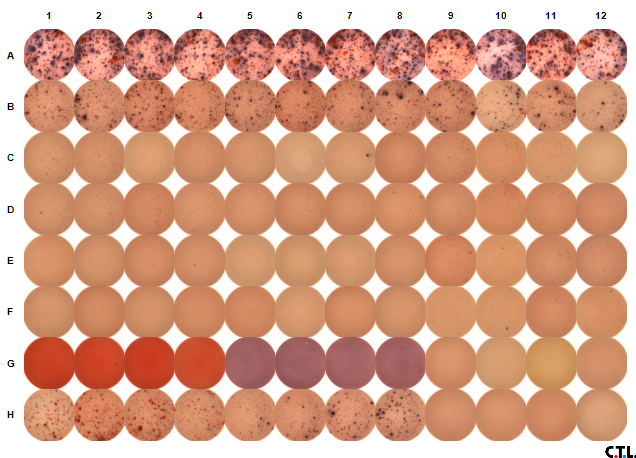

**B**

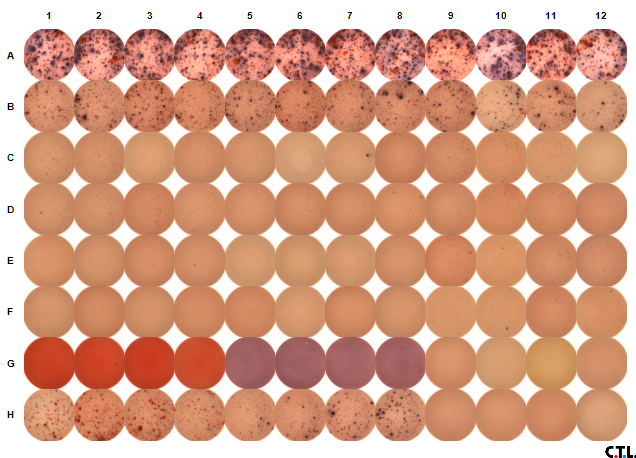

**C**

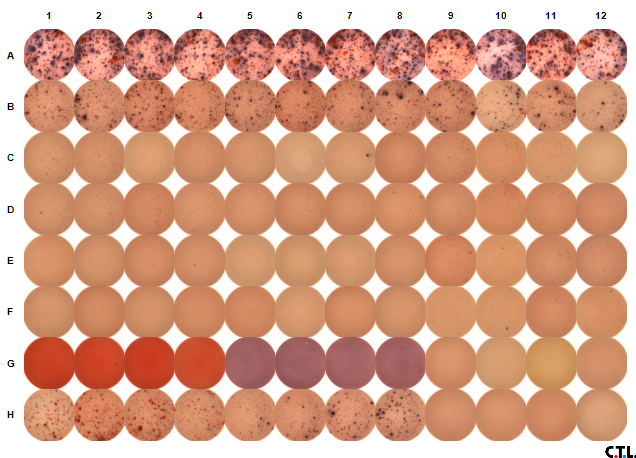

**D**

**IL-17**

**IFN-γ**

**Media Only**

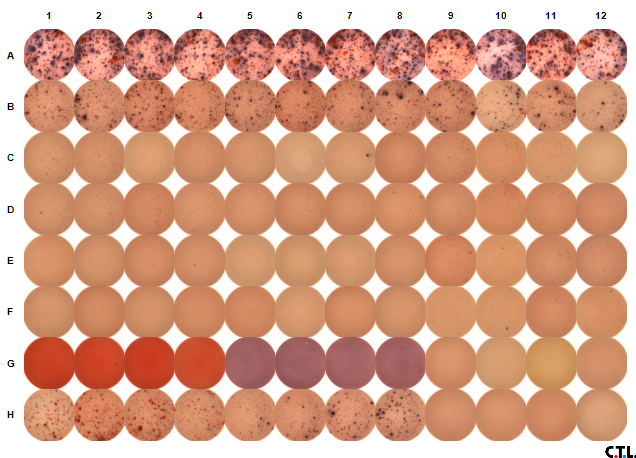

**E**

**Figure S14**. ELISPOT membranes from analysis of the TF-MUC4-B13G and TF-MUC4-CRM197 vaccinations. All wells contained 2.3x10^5^ CD4^+^ and 2.0x10^5^ Dendritic Cells. Rows are from vaccinations with (A) TF-MUC4-β-1,3-Glucan AuNP, (B) TF-MUC4-CRM197 conjugate, (C) B13G AuNP and (D) PBS. Row (E) is a control row where 1x10^5^ CD4^+^ cells were stimulated with Phytohemagglutinin and stained for IL-17, IFN-γ (positive controls) and media only (negative control)

**^1^H NMR spectrum of OVA 21-mer peptide (without N-terminal PEG linker)**

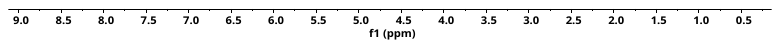

**^1^H NMR spectrum of OVA 21-mer peptide with N-terminal PEG linker**

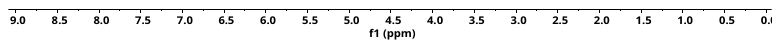

**Analytical HPLC of OVA 21-mer peptide with N-terminal PEG linker**

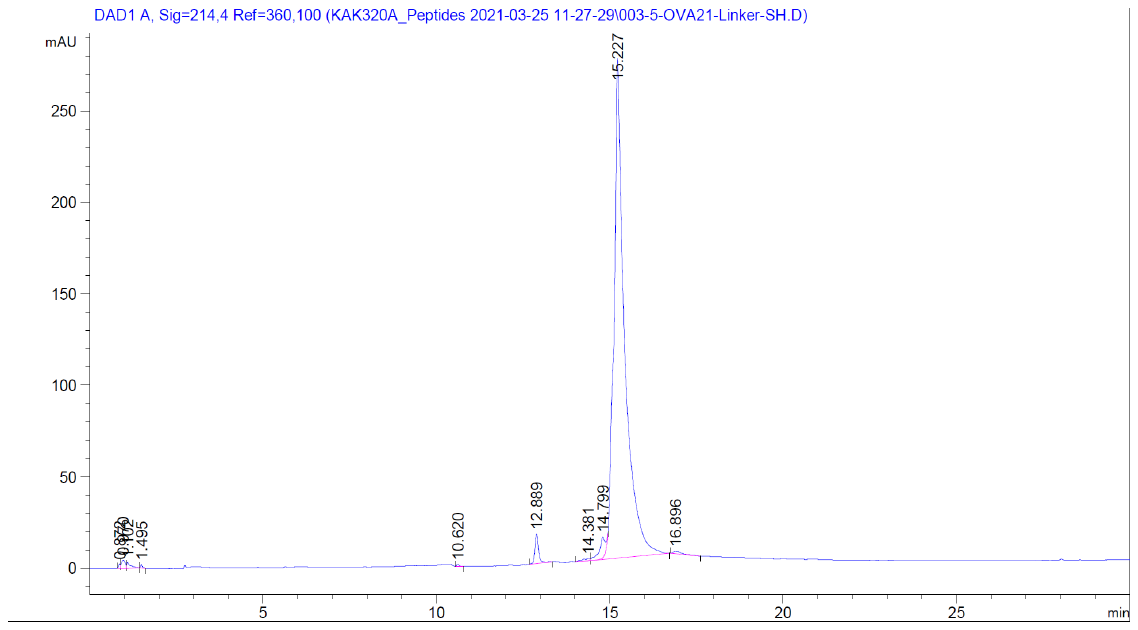

**High Resolution MS: OVA 21-mer peptide with N-terminal PEG linker**

Measured [M+ 3H]3+ =	m/z 895.1364 (Δ = 2.8 ppm)

Calculated [M+ 3H]3+ =	m/z 985.1339 (for C117H194N33O37S; z = 3)

Measured [M+ 2H]2+ =	m/z 1342.2013 (Δ = 3.1 ppm)

Calculated [M+ 2H]2+ =	m/z 1342.1972 (for C117H193N33O37S; z = 2)

Xtract -	Deconvoluted MW =	m/z 2682.3875 (Δ = 2.8 ppm)

Calculated MW =	m/z 2682.3799 (for C117H191N33O37S)

**Analytical HPLC of PEG-(GFLG)-MUC4 20 mer**

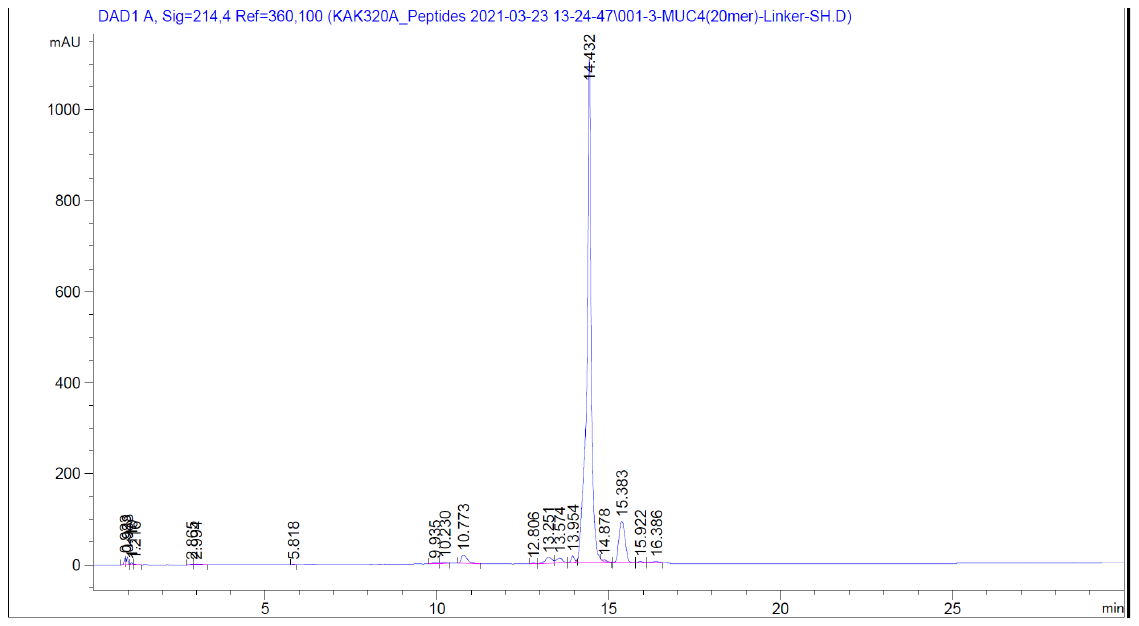

**^1^H NMR of PEG-(GFLG)-MUC4 20 mer**

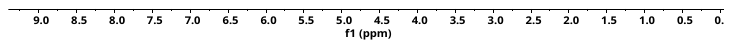

**^13^C NMR of PEG-(GFLG)-MUC4 20 mer** (Arrows indicate **C**F_3_**C**OOH carbons)

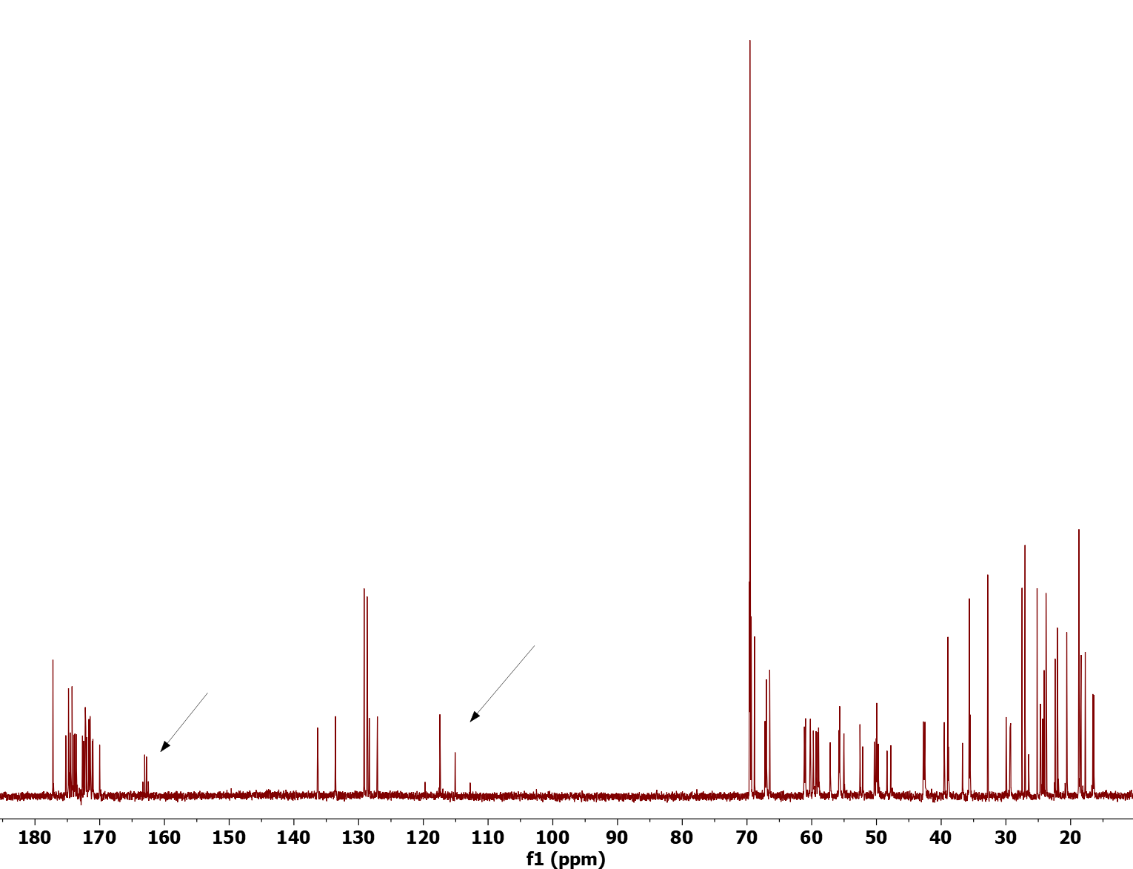

**High Resolution MS: PEG-(GFLG)-MUC4 20 mer**

Measured [M+ 3H]3+ =	m/z 817.7506 (Δ = 2.0 ppm)

Calculated [M+ 3H]3+ =	m/z 817.7490 (for C107H178N25O38S; z = 3)

Measured [M+ 2H]2+ =	m/z 1226.1223 (Δ = 2.0 ppm)

Calculated [M+ 2H]2+ =	m/z 1226.1198 (for C107H177N25O38S; z = 2)

Xtract -	Deconvoluted MW =	*m/z* 2450.230*3* (Δ = 2.2 ppm)

Calculated MW =	*m/z* 2450.2250 (for C_107_H_175_N_25_O_38_S)

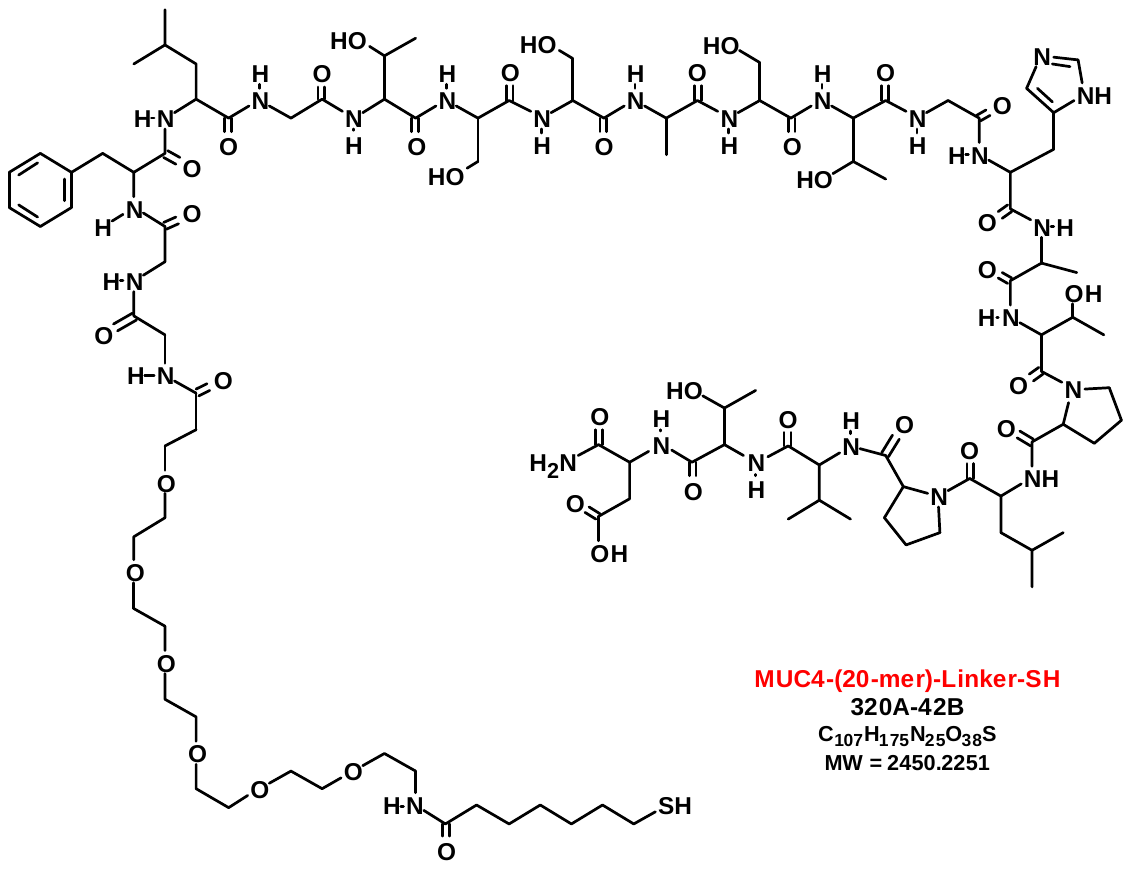

**Analytical HPLC of PEG-GFLG-TF-MUC4 20-mer**

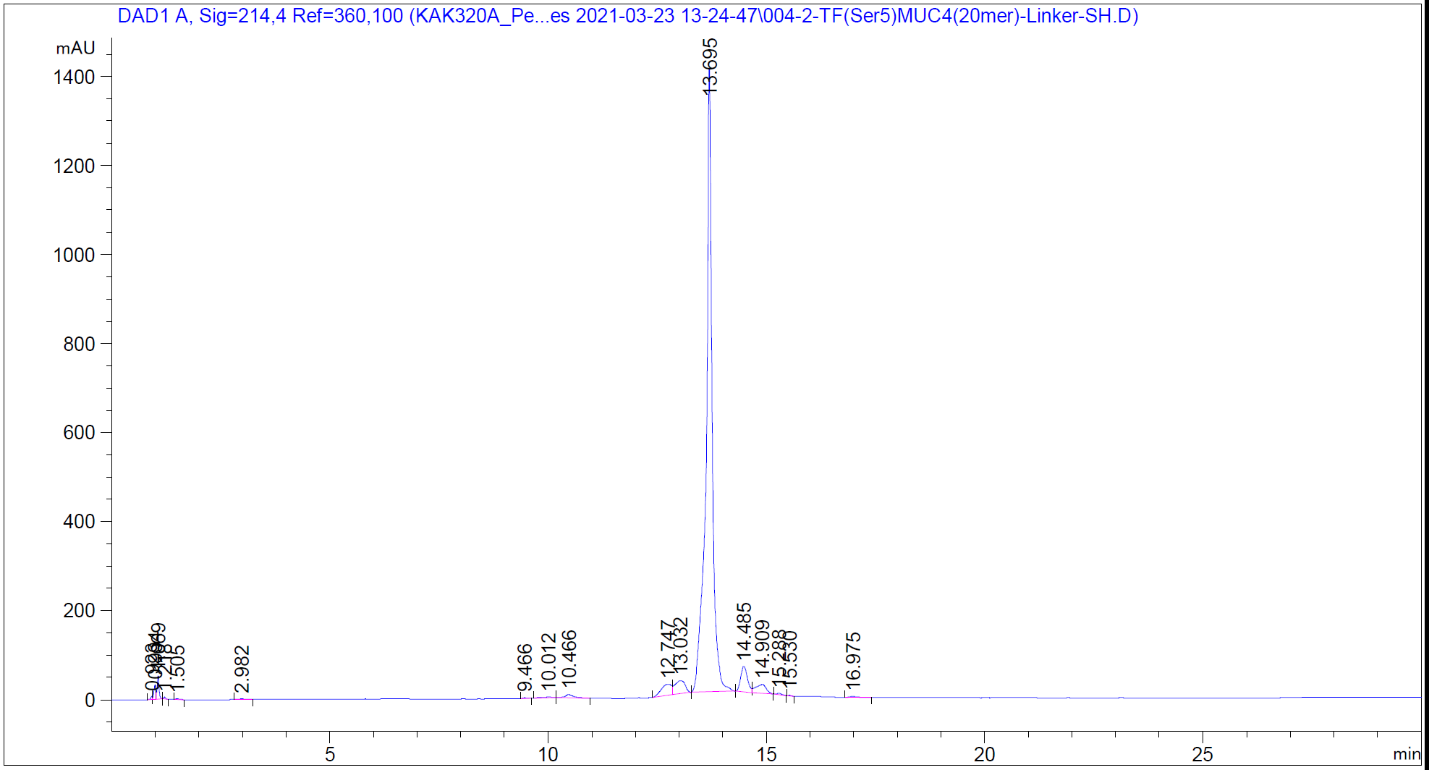

**^1^H NMR of PEG-(GFLG)-TF-MUC4 20 mer**

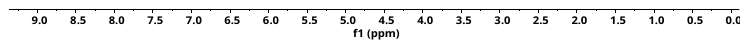

**^13^C NMR of PEG-(GFLG)-TF-MUC4 20 mer** (Arrows indicate **C**F_3_**C**OOH carbons)

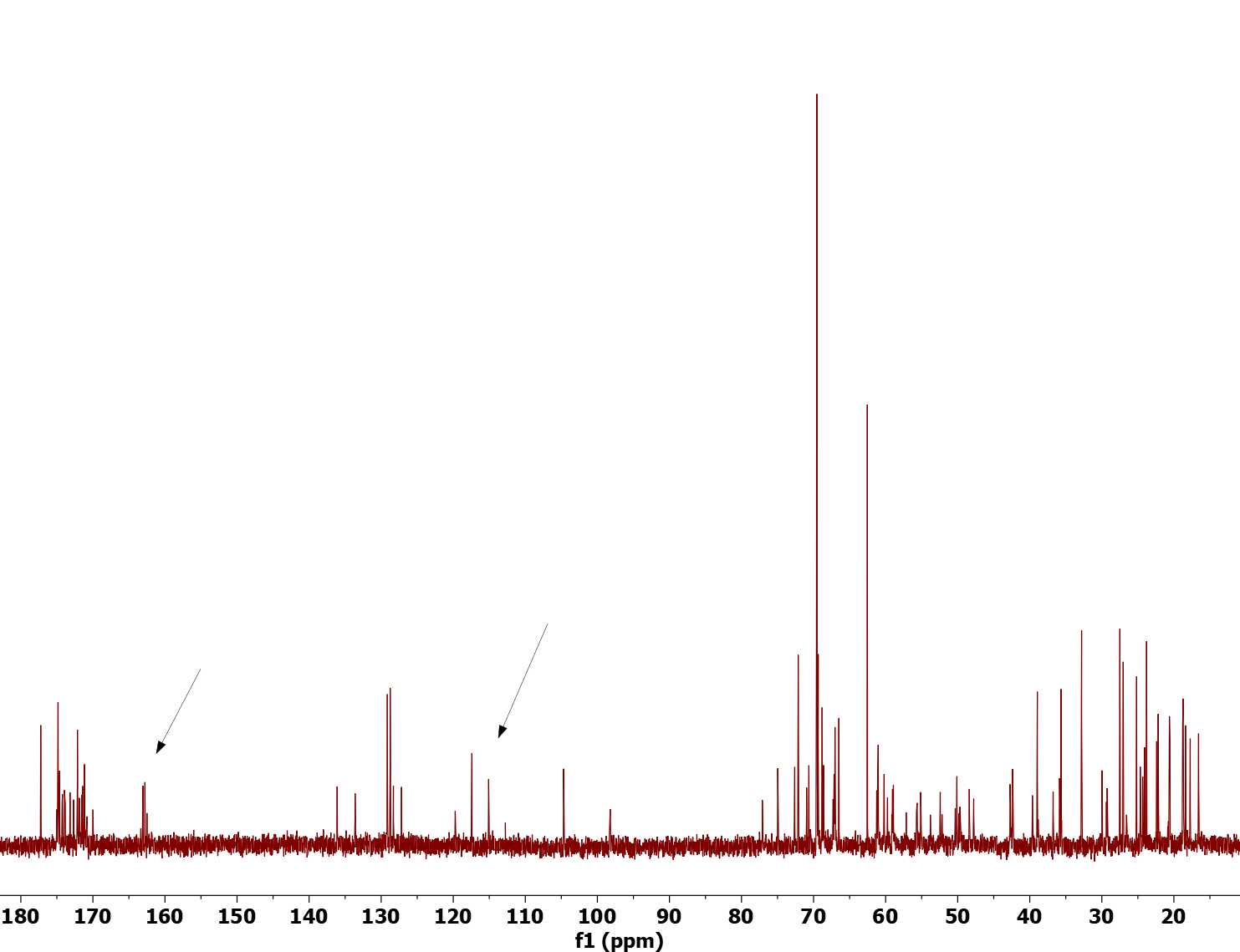

**High Res MS: PEG-GFLG-TF-MUC4 20-mer**

Measured [M+ 3H]^3+^ =	*m/z* 939.4627 (Δ = 3.2 ppm)

Calculated [M+ 3H]^3+^ =	*m/z* 939.4597 (for C_121_H_201_N_26_O_48_S; *z* = 3)

Measured [M+ 2H]^2+^ =	*m/z* 1408.690*5* (Δ = 3.3 ppm)

Calculated [M+ 2H]^2+^ =	*m/z* 1408.6859 (for C_121_H_200_N_26_O_48_S; *z* = 2)

Xtract -	Deconvoluted MW =	*m/z* 2815.366*4* (Δ = 3.3 ppm)

Calculated MW =	*m/z* 2815.3572 (for C_121_H_198_N_26_O_48_S)

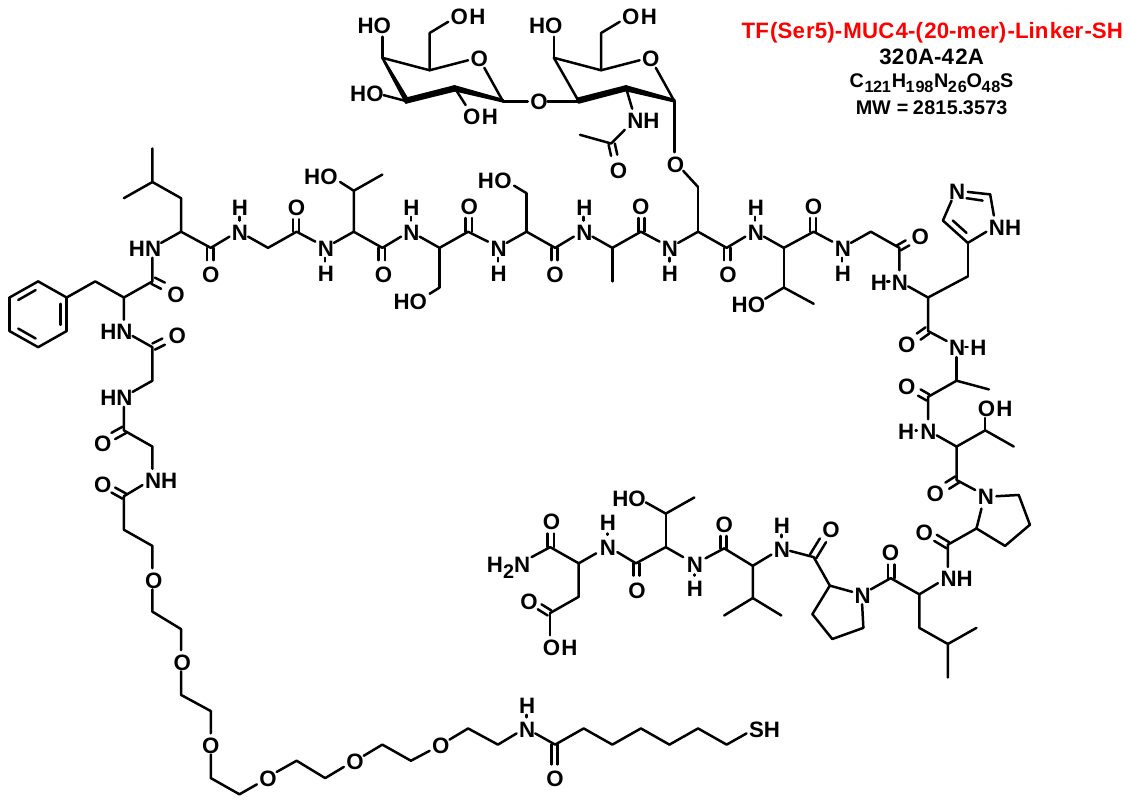
